# Target deconvolution of an insulin hypersecretion-inducer acting through VDAC1 with a distinct transcriptomic signature in beta-cells

**DOI:** 10.1101/2024.12.26.629558

**Authors:** Gitanjali Roy, Andrea Ordóñez, Derk D. Binns, Karina Rodrigues dos Santos, Michael B. Kwakye, George C. King, Noyonika Mukherjee, Andrew T. Templin, Zhiyong Tan, Timothy I. Richardson, Emma H. Doud, Amber L. Mosley, Christopher H. Emfinger, Alan D. Attie, Mark P. Keller, Travis S. Johnson, Michael A. Kalwat

## Abstract

Obesity, insulin resistance, and a host of environmental and genetic factors can drive hyperglycemia, causing β-cells to compensate by increasing insulin production and secretion. In type 2 diabetes (T2D), β-cells under these conditions eventually fail. Rare β-cell diseases like congenital hyperinsulinism (HI) also cause inappropriate insulin secretion, and some HI patients develop diabetes. However, the mechanisms of insulin hypersecretion and how it causes β-cell dysfunction are not fully understood. We previously discovered small molecules (e.g. SW016789) that cause insulin hypersecretion and lead to a loss in β-cell function without cell death. Here, we uncover the protein target of SW016789 and provide the first time-course transcriptomic analysis of hypersecretory responses versus thapsigargin-mediated ER stress in β-cells. In mouse MIN6 and human EndoC-βH1 β-cells, we identified and validated VDAC1 as a SW016789 target using photoaffinity proteomics, cellular thermal shift assays, siRNA, and small molecule inhibitors.

SW016789 raises membrane potential to enhance Ca^2+^ influx, potentially through VDAC1. Chronically elevated intracellular Ca^2+^ appears to underpin the negative impacts of hypersecretion, as nifedipine protected against each small molecule hypersecretion inducer we tested. Using time- course RNAseq, we discovered that hypersecretion induced a distinct transcriptional pattern compared to ER stress. Clustering analyses led us to focus on ER-associated degradation (ERAD) as a potential mediator of the adaptive response. SW016789 reduced the abundance of ERAD substrate OS-9 and pharmacological inhibition of ERAD worsened β-cell survival in response to hypersecretory stress. Changes in other ERAD components in MIN6 and EndoC-βH1 at the protein level were minor with either SW016789 or thapsigargin. However, immunostaining for core ERAD components SEL1L, HRD1, and DERL3 in non-diabetic and T2D human pancreas revealed altered distributions of SEL1L/HRD1 and SEL1L/DERL3 rations in β-cells of T2D islets, in alignment with altered ERAD in stressed β-cells. We conclude that hypersecretory stimuli, including SW016789- mediated VDAC1 activation, cause enhanced Ca^2+^ influx and insulin release. Subsequent differential gene expression represents a β-cell hypersecretory response signature that is reflected at the protein level for some, but not all genes. A better understanding of how β-cells induce hypersecretion and the mechanisms of negative feedback on secretory rate may lead to the discovery of novel therapeutic targets for T2D and HI.

## INTRODUCTION

Pancreatic islet β-cells must appropriately secrete insulin to maintain glucose homeostasis. Failures in this process lead to type 2 diabetes (T2D), which afflicts over 500 million people worldwide ^1^. In T2D, impaired production and response to insulin leads to chronic hyperglycemia and cardiovascular comorbidities. β-cells normally respond to increased concentrations of circulating nutrients by secreting insulin, which travels to peripheral tissues to stimulate glucose clearance ^2^. In healthy adults, fasting blood glucose concentration is ∼5 mM which can rise to 10-15 mM after a meal ^2^. β-cells sense this increase through passive glucose uptake and subsequent glycolytic and mitochondrial metabolism ^3^. These metabolic events cause oscillations in [ATP] and [ADP], with ATP stimulating closure and ADP stimulating opening of ATP-sensitive potassium channels (K_ATP_) ^4^. K_ATP_ closure leads to membrane depolarization, voltage-dependent Ca^2+^ channel (VDCC) opening, and Ca^2+^ influx, which triggers insulin exocytosis ^5–11^. Concurrently, metabolic amplification augments exocytosis without further Ca^2+^ influx, accounting for ∼50% of secreted insulin ^12–14^. While insulin secretion is necessary for glucose homeostasis, this process can go awry when secretory rates are elevated inappropriately in rare diseases like congenital hyperinsulinism (HI), in which insulin hypersecretion occurs. Insulin hypersecretion has also been postulated to contribute to prediabetes, T2D, and some cases of monogenic diabetes via multiple pathways, including oxidative stress and lipid signaling ^15^, palmitoylation ^16^, and genetic mutations ^17, 18^. We define the hypersecretory response as β-cell dysfunction occurring due to an abnormally high secretory rate caused by pharmacological, environmental, or genetic factors. Insulin hypersecretion can occur before the development of insulin resistance ^19, 20^. However, the mechanisms behind insulin hypersecretion and the associated responses remain incompletely understood. It is also unknown to what extent insulin hypersecretion may instigate β-cell failure and which steps might be therapeutically targetable. In some cases, diabetes treatments that promote secretion have been shown to exacerbate β-cell dysfunction ^21–23^. The hypersecretion response can be reversible ^21, 24^, which means interventions could lead to recovery of function. To prevent β-cell dysfunction in T2D and other genetic diseases, therapies are needed to alleviate hypersecretion while maintaining glucose homeostasis.

Hyperglycemia can also drive β-cell failure independent of insulin hypersecretion^25^, further supporting the need for multiple treatment strategies.

Insulin hypersecretion may present as repeated occurrences for short durations, or chronically elevated secretory rates, and is known to be induced by multiple small molecules and drugs ^21, 24, 26–28^. In response to these types of treatments, we and others have found β-cells adapt by shutting down secretory function, perhaps as a mechanism to avoid cell death ^21, 24, 26^. The mechanisms of pharmacologically induced insulin hypersecretion and its downstream impacts on β-cell function are not fully understood. We previously identified a small molecule, SW016789, which causes insulin hypersecretion in acute stimulations (1-2 h) and results in a loss in β-cell function after longer (4-24 h) exposures ^24^. We determined that SW016789 enhances nutrient-stimulated Ca^2+^ influx to elicit hypersecretion, subsequently causing loss of stimulated secretion and a transient ER stress-like response, but without inducing cell death. The proximal mechanism of action for SW016789 and the nature of the β-cell hypersecretory response were unclear.

In the current study, we report the discovery of a candidate target protein for SW016789 action, the voltage-dependent anion channel VDAC1. Further, we find that hypersecretion can occur in response to structurally diverse compounds and that the key steps likely lie downstream of enhanced Ca^2+^ influx. To better understand the processes involved in the hypersecretory response, we used time course transcriptomics and clustering analyses in β-cells exposed to SW016789 compared to thapsigargin treatment. These data elucidated a unique hypersecretory response signature. This signature is enriched for multiple pathways, including core components of ER-associated degradation (ERAD), COP-II transport, serine and one-carbon metabolism, and certain ER chaperones. We found that ERAD may play a part in the β-cell adaptive response to hypersecretory stress to avoid cell death and that the ratio between ERAD components may be altered in islets in T2D.

## RESULTS

### SW016789 induces hypersecretory stress in human β-cells and interacts with voltage-dependent anion channel (VDAC1)

We previously published the phenotypic effects of SW016789-induced hypersecretory stress in β-cells ^24^. However, the exact pathways through which hypersecretion acts are not well understood. Using an XBP1 splicing-based reporter, hypersecretion induced by SW016789 caused a transient and relatively low-amplitude ER stress response in human EndoC-βH1 β-cells compared to the sustained response elicited by thapsigargin (**Fig 1A**).

**Figure 1.**
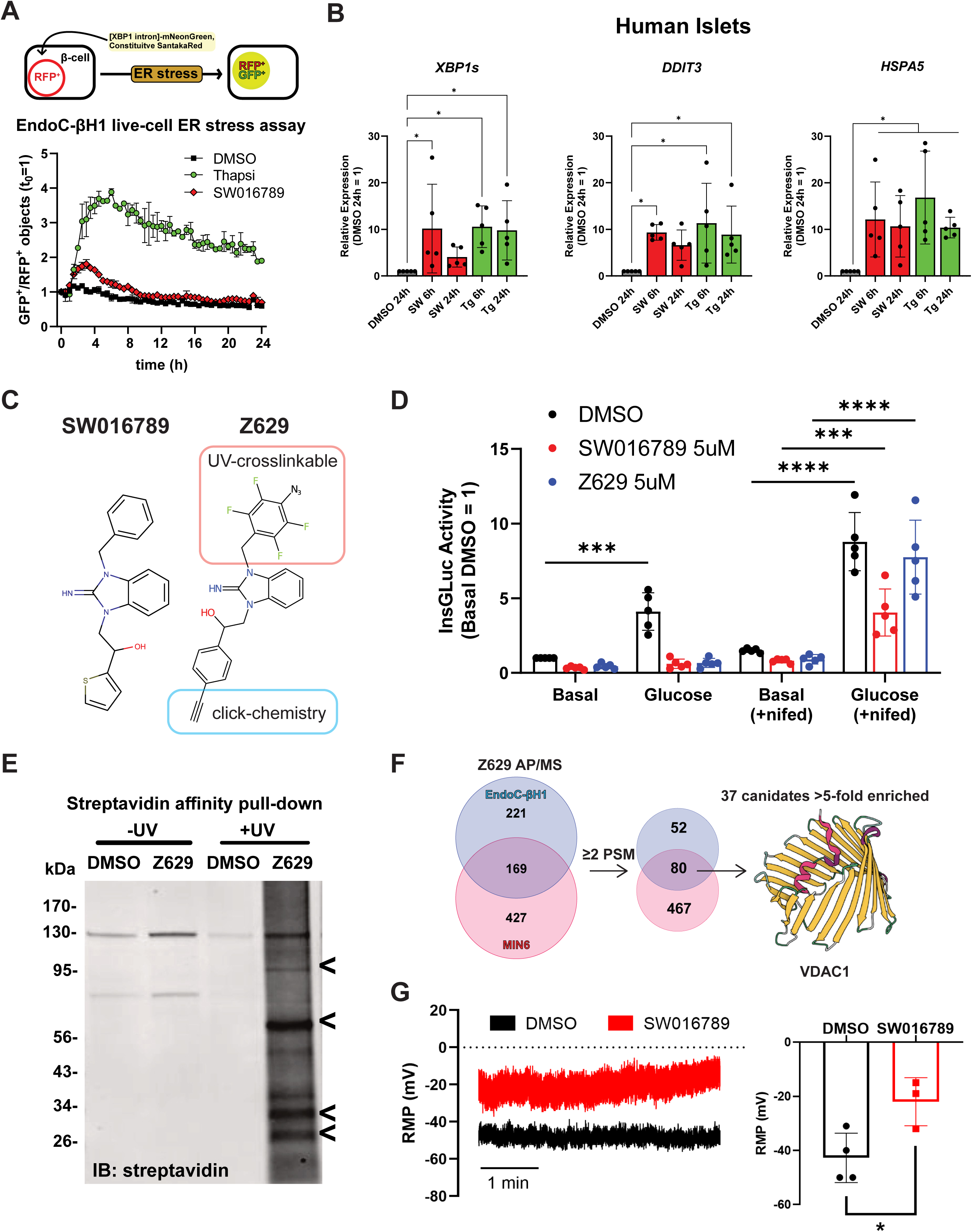
VDAC1 is implicated in hypersecretion responses induced by SW016789 in human and mouse β-cells. **A)** SW016789 (10 μM) induces a transient response in human EndoC-β1 cells expressing a fluorescent XBP1 splicing-based ER stress sensor, while thapsigargin (3 μM) induced a sustained response. **B)** Gene expression measured by qPCR in human islets treated with SW016789 (SW, 10μM), thapsigargin (Tg, 1 μM), or vehicle (DMSO, 0.1%) for 6 or 24 h. Data are the mean ± SD of N=5. *, P<0.05. **C)** Chemical structure of SW016789 and the photoaffinity probe Z629. **D)** InsGLuc secretion assay in β-cells treated 24 h with DMSO, SW016789 (5 μM), or Z629 (5 μM) in the presence or absence of nifedipine (10 μM). Data are the mean ± SD of N = 5. *, P<0.05. **E)** Streptavidin western blot indicating biotin-conjugated proteins in MIN6 cells UV-crosslinked to Z629, followed by click-medicated reaction with biotin-azide. **F)** Mass spectrometry-based chemical proteomics was used to identify VDAC1 as a target of SW016789 in MIN6 and EndoC-βH1 cells. **G)** Resting membrane potential measurements in MIN6 cells treated with DMSO (0.1%) or SW016789 (5 μM). Data are the mean ± SD of N ≥ 3. *, P<0.05.

The transient nature of hypersecretory stress from SW016789 compared to sustained ER stress from thapsigargin was consistent in isolated human islets treated for either 6 or 24h, as shown by spliced *XBP1* and *DDIT3* (CHOP) expression (**Fig 1B**). However, *HSPA5* (BiP) expression was sustained with both treatments. From our previous findings, it was clear that SW016789-induced hypersecretory stress depended on Ca^2+^ influx, but the exact protein target(s) were unknown. To address this, we leveraged structure-activity-relationship data using a small set of chemical analogs of SW016789 ^24^. Based on the activity of those analogs, we determined that the central hydroxyl group could not be modified and that adding side chains (e.g. azide or alkyne) may be tolerated on the benzyl and thiophene rings. We designed a photoaffinity probe, Z629, to perform direct target identification experiments **(Fig 1C, S1A).** To verify that Z629 retained activity, we treated InsGLuc- MIN6 cells for 24 h in the presence of DMSO, SW016789, or Z629. We observed that both SW016789 and Z629 suppressed glucose-stimulated secretion (**Fig 1D**). We previously showed that the dihydropyridine voltage-dependent Ca^2+^ channel (VDCC) blocker nifedipine protected against the inhibitory effects of SW016789 ^24^. Using this paradigm, we observed that nifedipine also protected β-cells against Z629 (**Fig 1D**). These data suggested that Z629 has similar activity to the parental SW016789 compound. The Z629 probe incorporates an alkyne for click chemistry and a tetrafluorophenyl azide for UV-crosslinking. Upon UV exposure, the azide forms a reactive nitrene group that forms covalent bonds with nearby proteins. Alkyne-labeled proteins can be modified with biotin via click chemistry, followed by affinity purification with streptavidin-agarose and analysis by mass spectrometry. Having established SW016789 activity in mouse and human β-cell lines, we performed target identification studies in both MIN6 and EndoC-βH1 β-cells to increase the chances of identifying relevant targets. We treated both lines with Z629 or DMSO and exposed the cells to UV-B light. The lysates were subjected to click chemistry and affinity purification followed by immunoblotting to confirm biotin labeling (**Fig 1E**). Bands present in the non-UV-treated samples are endogenously biotinylated proteins (e.g. pyruvate carboxylase, ∼130kDa). The experiment was repeated, and samples were subjected to on-bead digestion for proteomic analysis. We identified 547 potential Z629-interacting proteins in MIN6 and 132 proteins in EndoC-βH1 β-cells with ≥2 spectral counts in the Z629-treated samples (**Fig 1F, Table S2**). Intersecting the MIN6 and EndoC- βH1 lists and applying a >5-fold enrichment cutoff narrowed this list to 37 candidates. The voltage-dependent anion channels, VDAC1, VDAC2, and VDAC3, were detected in Z629 pulldowns from both MIN6 and EndoC-βH1 cells, consistent with their expression in both cell lines and in human islets (**Fig S1B**). VDAC1 stood out among the candidates because it has been implicated in the β- cell in T2D and has the potential to alter membrane potential which may explain the effects of SW016789 ^29^. Supporting this, we observed increased resting membrane potential in MIN6 cells exposed to SW016789 (**Fig 1G**).

### VDAC1 is a direct target of SW016789 and is necessary for full SW016789 activity in β-cells

To validate VDAC1as a target we applied three orthogonal methods. First, we depleted *Vdac1* in InsGLuc-MIN6 β-cells using siRNA (**Fig 2A**). *Vdac1*-depleted cells had a significantly reduced response to acute SW016789-enhanced glucose-stimulated secretion (**Fig 2B**). However, KCl-stimulated secretion was intact in *Vdac1*-depleted cells (**Fig 2C**), suggesting that membrane depolarization and secretion induced solely by Ca^2+^ influx were not impaired. Second, we co-treated cells for 24 hours with SW016789 alone or in combination with the VDAC1 inhibitor VBIT4 or nifedipine. Treatment with SW016789 alone inhibited subsequent glucose-stimulated secretion, as expected (**Fig 2D**) ^24^. However, co-treatment of SW016789 with either VBIT4 or nifedipine protected against this inhibition (**Fig 2D**). Third, we applied the cellular thermal shift assay (CETSA), which uses a temperature gradient to assess alterations in the melting temperature of the protein of interest upon binding of a ligand ^30^. We exposed MIN6 β-cells to a temperature gradient of 37-75°C to determine endogenous VDAC1 aggregation in the absence of ligand. Immunoblotting shows the stability of Vdac1 at temperatures up to 71°C (**Fig S2A**). Next, we shifted to a higher temperature gradient (55-80°C) in the presence of DMSO or SW016789 (10 μM). SW016789 caused a significant shift in VDAC1stability at 59.9 to 64.3°C compared to DMSO (**Fig 2E**). Isothermal concentration dependence of SW016789 (0-20 μM) on Vdac1 stability was then performed at 64.3°C in MIN6 cells. VDAC1stability appeared to increase in response to increasing concentrations of SW016789 and was increased significantly at 20 μM SW016789 (**Fig 2F**). Based on these data, we conclude that SW016789 can act through VDAC1 to cause hypersecretion, possibly via increasing membrane ion permeability and raising membrane potential, thereby potentiating glucose-stimulated Ca^2+^ influx (**Fig 2G**).

**Figure 2.**
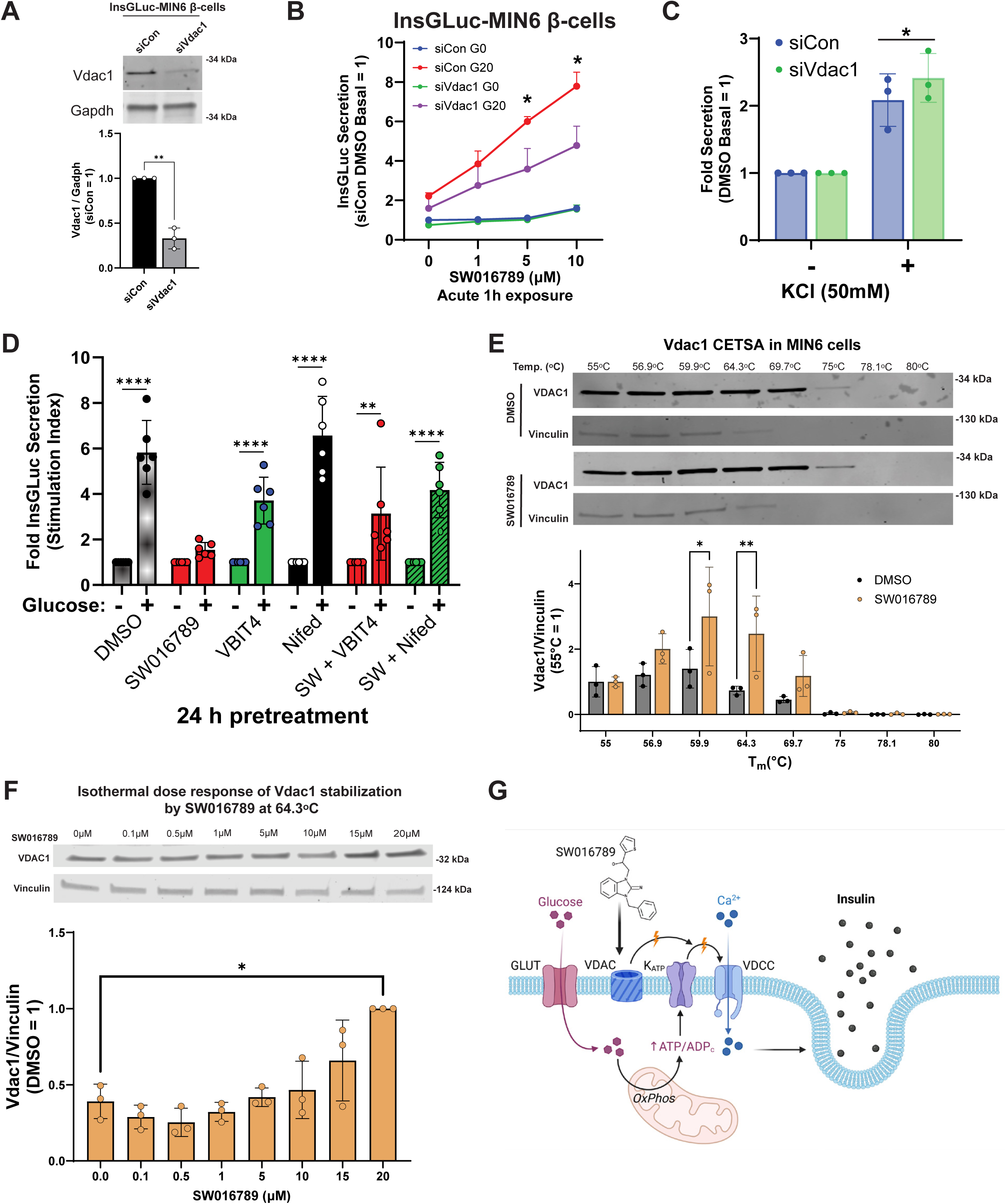
VDAC1 is stabilized by SW016789 and required for SW016789 activity. **A)** siRNA knockdown of *Vdac1* in InsGLuc-MIN6 β-cells. Data are the mean ± SD of N=3. *, P<0.05. **B)** VDAC1 is required in InsGLuc-MIN6 cells for full secretory response to SW016789 in the presence of 20 mM glucose. **C)** *Vdac1* knockdown does not affect KCl-induced secretion. **D)** 24 h co- treatment with VDAC1 inhibitor VBIT4 (10 μM) or nifedipine (10 μM) blocks the SW016789-induced loss of secretory response. **E)** Cellular thermal shift assay (CETSA) in MIN6 β-cells shows SW016789 binds and stabilizes VDAC1 significantly at 64.3°C compared to DMSO. Data are the mean ± SD of N=3. *, P<0.05. **F)** Isothermal dose-response CETSA shows VDAC1 stabilization is significant in the presence of 20 μM SW016789 MIN6 β-cells. Data are the mean ± SD of N=3. *, P<0.05. **G)** Working model of SW016789-mediated VDAC1 activation, membrane depolarization, and enhanced Ca^2+^ influx and hypersecretion.

### Small molecule-mediated hypersecretion-induced loss of β-cell function requires chronic Ca^2+^ influx

Many physiological and pharmacological stimuli can enhance Ca^2+^ influx and potentially cause hypersecretion, likely through multiple mechanisms including VDAC1. We previously showed that ≤2 h exposure to SW016789 potentiated nutrient-induced Ca^2+^ influx to cause insulin hypersecretion ^24^. However, ≥4 h of exposure significantly inhibited GSIS responses without causing cell death. We hypothesized that the effects of SW016789 may not be unique, as other pharmacological stimulators of insulin secretion have been shown to lead to suppression of β-cell function, as is the case for the NMDA receptor antagonist dextrorphan (DXO) ^26^ and sulfonylureas ^21, 24, 26, 27, 31^. Consistently, we observed a loss of function in MIN6 cells treated chronically with DXO or the sulfonylurea glimepiride (**Fig 3A**). We also discovered structurally distinct small molecule hypersecretion inducers in a previous high-throughput screen (**Fig S3A**) ^32^. These compounds had similar activity to SW016789, with 24h pre-treatment leading to suppressed Ca^2+^ influx (**Fig S3B-D**) and insulin secretion responses (**Fig 3B**) in MIN6 cells. In each of these cases, preventing Ca^2+^ influx using the VDCC blocker nifedipine conferred protection from the chronic inhibitory effects of these hypersecretion inducers (**Fig 3A,B**). However, nifedipine was not able to protect against thapsigargin-induced loss of function, distinguishing the hypersecretory response from ER stress. Considering the distal effects of hypersecretion on β-cell function, from here we used SW016789 as a hypersecretion-inducing tool compound to investigate the downstream dynamics of this response in β-cells and to compare with ER stress responses.

**Figure 3.**
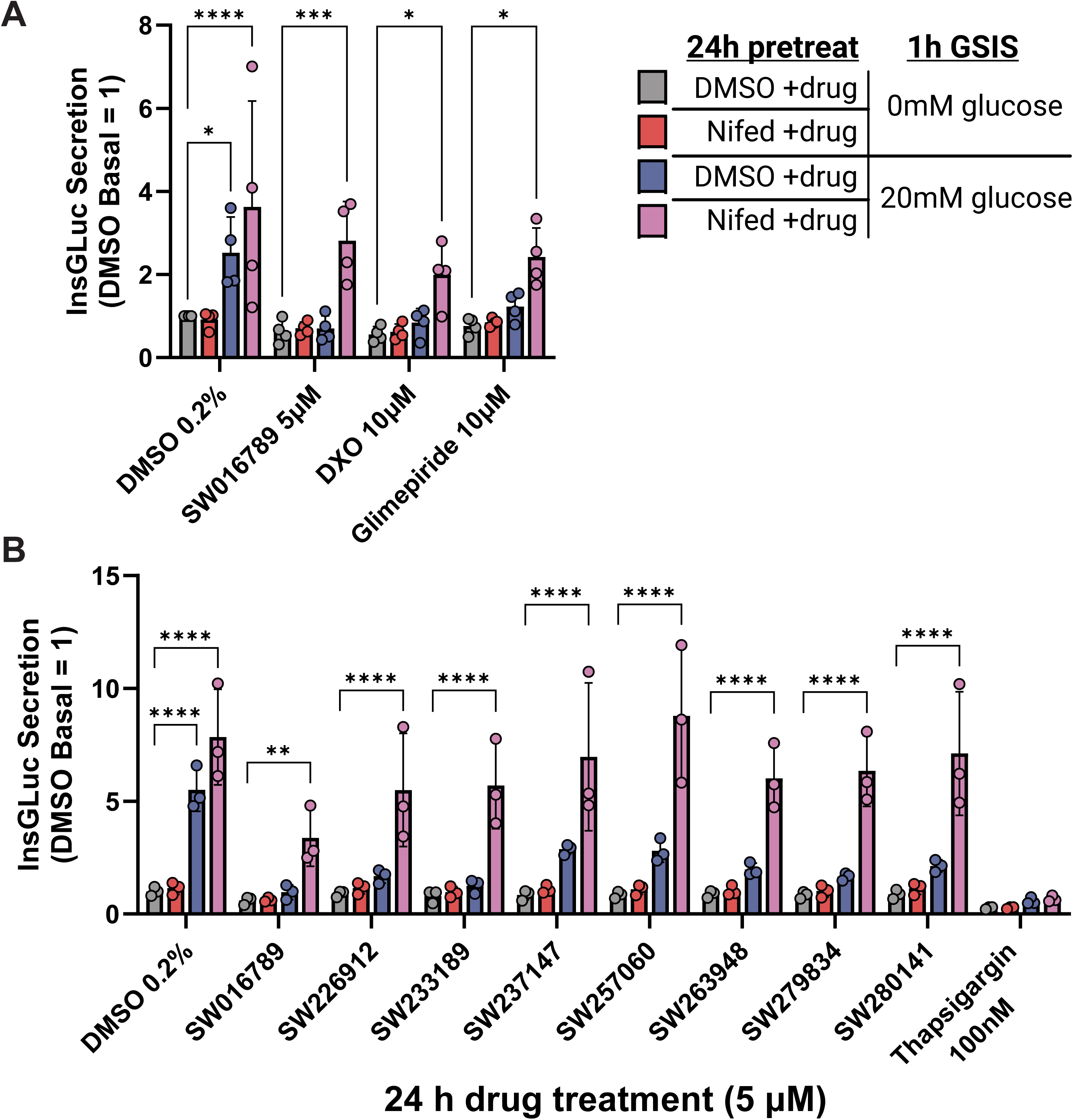
Ca^2+^ influx is central to the loss of β-cell function in response to structurally divergent small molecule hypersecretion inducers. **A)** InsGLuc-MIN6 cells were treated 24 h with indicated small molecules (DMSO, SW016789, dextrorphan (DXO), glimepiride) in the presence or absence of nifedipine (10 μM). Cells were then assessed for their glucose-stimulated secretion response in the absence of small molecules. Data are the mean ± SD of N=4. *, P<0.05. **B)** InsGLuc-MIN6 cells were treated as in (**A**), except structurally distinct hypersecretion-inducing compounds were tested (5 μM), as well as thapsigargin (100 nM). Data are the mean ± SD of N=3. *, P<0.05.

#### Identification and analysis of time-dependent transcriptomic changes in response to hypersecretory stress in β-cells

To unbiasedly identify transcriptomic alterations downstream of hypersecretory stress, MIN6 β-cells were treated with DMSO, SW016789, and thapsigargin for 1,2,6 and 24 hours and submitted for RNA-sequencing (**Fig 4A, Table S3**). Time course differential gene expression analysis in edgeR (|log_2_FC| > 1 & FDR < 0.05) indicated 926 genes altered by thapsigargin and 168 genes altered by SW016789 with 109 in common (**Fig 4B**). Given our preexisting knowledge of β-cell-relevant pathways and the effects of hypersecretion and ER stress, we noted distinct expression differences between SW016789 and thapsigargin for multiple pathways (**Fig 4C**). For example, thapsigargin altered expression genes involved in the unfolded protein response, ER-associated degradation (ERAD), redox stress, autophagy, cell death, insulin processing, and secretion, disallowed β-cell genes, metabolism, and β-cell transcription factors. On the other hand, SW016789 selectively impacted subsets of the UPR, immediate early response, and insulin secretion genes. Notably, SW016789 did not alter certain maladaptive responses like *Txnip*, *Qrich1*, or many of the cell death genes. However, many distinctions in gene expression between SW016789 and thapsigargin appeared to be related to the temporal expression pattern. Therefore, to better understand these dynamic expression changes, we clustered the data using two different approaches. We used weighted gene co-expression network analysis (WGCNA) to cluster all genes in the raw RNA-seq count matrix into modules based on their expression patterns, regardless of significance or fold change in edgeR (**Fig 5A, Fig S5, Table S4**). For example, the ‘royalblue’ and ‘turquoise’ WGCNA modules stood out as robustly changed in opposite directions by SW016789 and thapsigargin (**Fig 5B**). We performed gene set enrichment analysis on each module using EnrichR-KG and created a directed network comprised of all modules with significantly enriched terms (**Fig 5C**). This network illustrated the distinct differences between hypersecretion and thapsigargin-mediated ER stress and showed positive correlations between WGCNA modules like ‘royalblue’, which was enriched for secretory pathway genes and ‘black’, which was enriched for protein processing in the ER.

**Figure 4.**
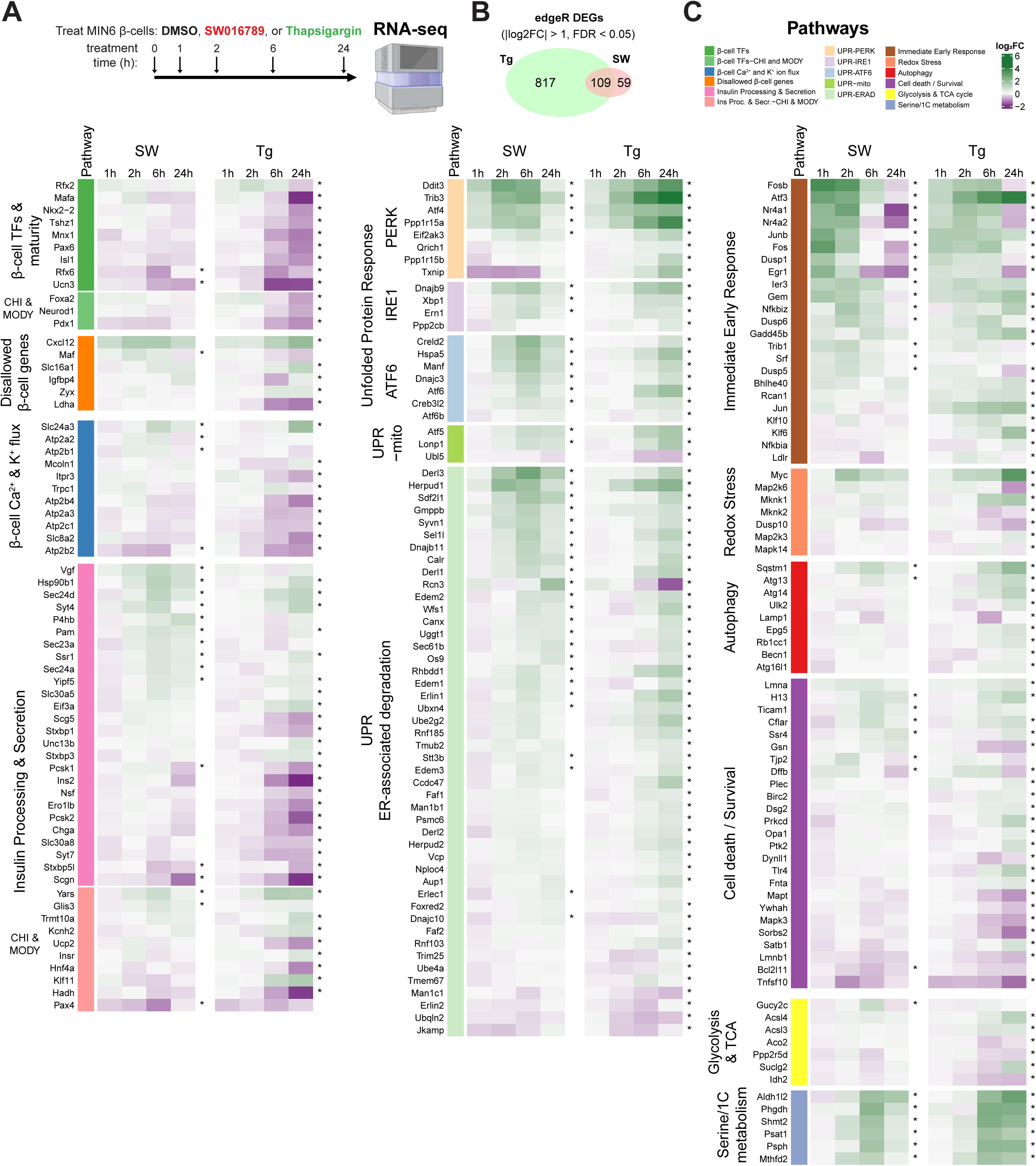
Time course temporal transcriptomics uncovers differential effects of hypersecretion and ER stress. **A)** MIN6 β-cells treated with DMSO (0.1%), SW016789 (SW, 5 μM), or thapsigargin (Tg, 100 nM) for 1, 2, 6 and 24 hours. N=3. **B)** Time course edgeR analysis results showing overlap of differentially-expressed genes (DEGs) between Tg and SW. DEGs are shown by log2 fold-change versus baseline. **C)** Heatmaps showing selected pathways containing DEGs that were significant in SW, Tg, or both. *, FDR<0.05 in at least one time point vs. baseline.

**Figure 5.**
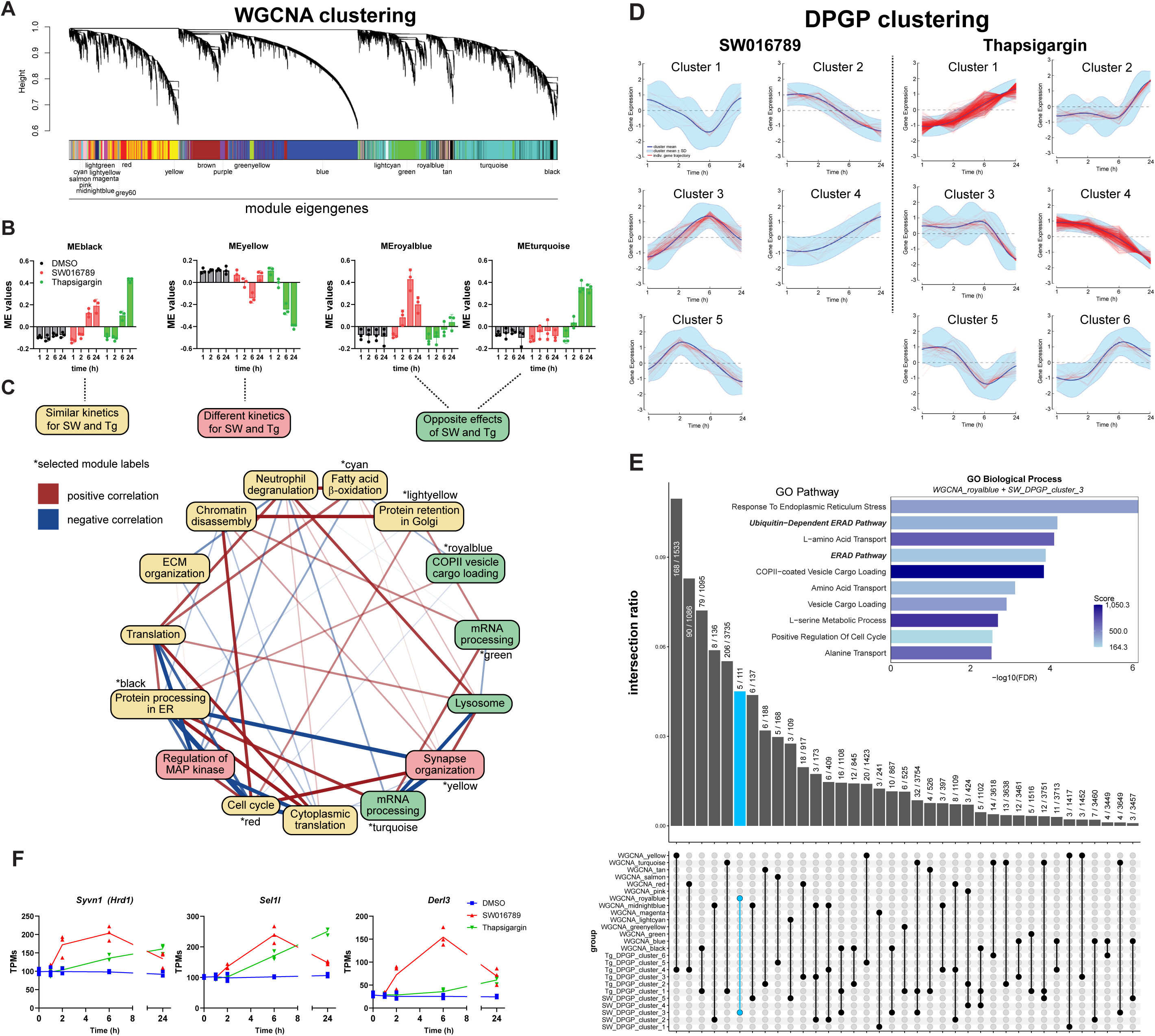
Clustering analyses unveil a distinct β-cell hypersecretory response signature. **A)** Weighted gene co-expression network analysis (WGCNA) of the entire unfiltered time course transcriptomics data set highlights multiple modules. **B)** Module eigengene (ME) values for modules that have similar kinetics between SW016789 (SW) and thapsigargin (Tg): black, different kinetics: yellow, or opposite directional effects: royalblue and turquoise. **C)** Directed network analysis of module eigengenes with their top enriched ontology shown and colored by their kinetics and effect directions. Edges connecting the nodes are colored by positive (red) or negative (blue) correlation, and thickness indicates the magnitude of the correlation, with thicker edges showing stronger relationships. **D)** Dirichlet process Gaussian process (DPGP) clustering analysis groups genes based on their temporal expression pattern. **E)** UpSet plot comparison of all WGCNA modules and DPGP clusters. The overlap between SW_DPGP_cluster_3 and ‘royalblue’ and ‘blue’ modules is highlighted in light blue. Because the expression dynamics of these two clusters matched well, the two lists were merged and analyzed by Enrichr. GO Biological Process enriched terms are shown in the inset. **F)** Select genes for the ERAD pathway show differential expression dynamics between SW and Tg. Values shown are TPMs from RNAseq data (N=3).

In parallel with WGCNA, we used Dirichlet process-Gaussian process (DPGP) based clustering of significant genes from **Fig 4A** based on the dynamics of their expression throughout the time course ^33^. Using the edgeR time course log_2_FC values as input, DPGP unbiasedly clustered the genes for SW016789 and thapsigargin (**Fig 5D**). Notably, most SW016789-altered genes fell into a single cluster (SW_DPGP_cluster_3), which followed a pattern of early downregulation, followed by increased expression peaking at 6 h and returning to baseline by 24 h. We also noted that SW_DPGP_cluster_2 and SW_DPGP_cluster_3 exhibited a pattern of peak expression at 1 and 2 h, respectively, likely due to the immediate early response downstream of enhanced Ca^2+^ influx. For thapsigargin, most genes fell into Tg_DPGP_cluster_1 and Tg_DPGP_cluster_2, which represent genes that continuously increase or decrease with time, respectively. We compared the WGNCA and DPGP clustering results using an UpSet plot ranked by the intersection ratio (**Fig 5E**). The UpSet plot indicated that the ‘royalblue’ WGCNA module was the most similar to SW_DPGP_cluster_3 but included additional genes that had not reached significance in edgeR analyses. SW_DPGP_cluster_3 also overlapped with Tg_DPGP_cluster_1, highlighting genes that are modulated transiently after hypersecretion but which are induced more slowly by ER stress. For our downstream analyses, we combined the gene lists of ‘royalblue’ and SW_DPGP_cluster_3 because genes falling into these dynamic expression patterns were similar. We propose this set represents a hypersecretion response signature. Within this signature, we found enrichment of pathways related to ERAD and ER stress (e.g. *Derl3*, *Syvn1, Sel1l, Hspa5, Ppp1r15a*), amino acid transport, vesicle trafficking (e.g. COP-II complex), and serine metabolism (e.g. *Psat1*, *Psph*, *Phgdh*) (**Fig 5E, inset**). ERAD has been implicated as an important pathway in β-cell function ^34–37^, and thus, we hypothesized it could serve an adaptive role during hypersecretion. ERAD mediates the extrusion and ubiquitination of misfolded proteins in the ER membrane and lumen, and thereby mitigates ER stress and eventual apoptosis. ERAD factors like *Sel1l, Derl3,* and *Syvn1* are induced by SW016789 at 6h, while their induction by Tg is either weaker or delayed (**Fig 5F**). We next investigated the involvement of ERAD in the β-cell hypersecretion response and diabetes pathology using pharmacological treatments, immunoblotting in murine and human β-cell lines, and immunostaining of human pancreas sections.

#### ERAD activity may be altered by hypersecretion and protect against apoptosis

As one readout of ERAD activity, we monitored the protein abundance of the ERAD substrate OS-9 ^38^. We treated β-cells with Tg, SW016789, and eeyarestatin I (ES1) which inhibits VCP/p97, an AAA+ ATPase responsible for peptide extrusion through the ERAD complex. SW016789 alone or in combination with ES1 caused a reduction in OS-9 abundance (**Fig 6A**), potentially reflecting increased ERAD flux. It is worth noting that OS-9 mRNA expression is increased by both SW016789 and Tg at 24h (**Fig 4C**). ERAD induction may also contribute to β-cell survival during the hypersecretory response while Tg-treated cells undergo chronic ER stress and apoptosis. To test this, we exposed β-cells to either ES1 or SW016789 alone or in combination for 24 h. Only the combination of ES1 and SW016789 increased caspase 3/7 activity (**Fig 6B**), suggesting a requirement for ERAD activity for β-cell survival during hypersecretion. MIN6 β-cells retained glucose responsiveness after 24 h treatment with ES1, indicating ERAD inhibition under these conditions is not causing major defects in β-cell function (**Fig 6C**). As expected, SW016789 alone or in combination with ES1 caused a loss of function (**Fig 6C**).

**Figure 6.**
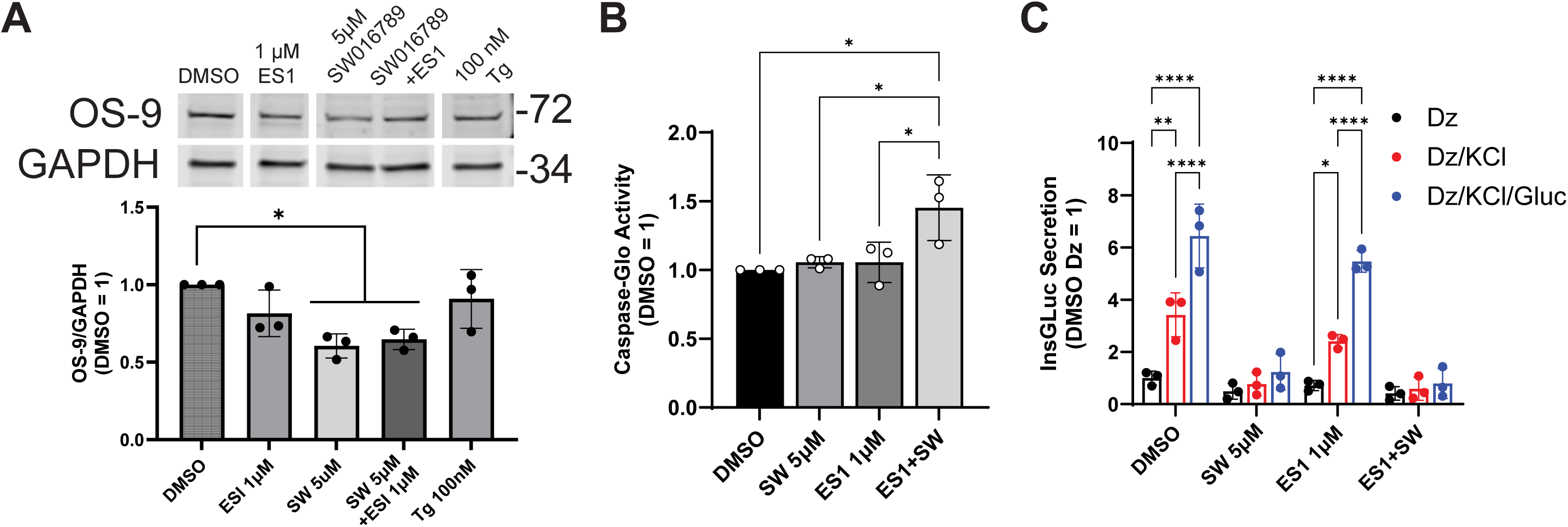
Regulation of ERAD is part of the transcriptomic signature of hypersecreting β- cells. **A)** MIN6 cells were treated for 24 h with 1 μM eeyarestatin I (ES1), 5 μM SW016789, ES1 + SW016789, or 100 nM thapsigargin (Tg). Expression of ERAD substrate OS-9 was monitored by immunoblotting. Data are the mean ± SD of N=3. *, P<0.05. **B)** Inhibiting ERAD with ES1 sensitizes β-cells to SW016789 hypersecretion-induced apoptotic ER stress. Data are the mean ± SD of N=3. *, P<0.05. **C)** 24 h treatment with 1 μM ES1 does not affect β-cell secretory response to glucose or to the chronic inhibitory effects of SW016789. Cells were stimulated acutely in the presence of diazoxide (250 μM), KCl (35 mM), and glucose (20 mM). Data are the mean ± SD of N=3. *, P<0.05.

#### Hypersecretion responses at the protein level induce the immediate early response and do not robustly engage canonical ER stress pathways

We next determined the extent to which transcriptional alterations due to hypersecretion or canonical ER stress led to changes in protein abundance. Mouse MIN6 cells and human EndoC-βH1 cells were treated with SW016789 or thapsigargin for 1, 2, 4, 6, and 24 h, as well as vehicle control DMSO for 24 h. In MIN6 β-cells treated with thapsigargin, we observed significantly increased amounts of phosphorylated eIF2α by 2h, followed by elevated CHOP, cleaved PARP, and BiP by 24 h (**Fig 7A-E**), However, with SW016789-induced hypersecretion, we observed a rapid elevation of cFos (**Fig 7H**) followed by increased PHGDH expression at 24 h (**Fig 7I**), with minimal impact on eIF2α phosphorylation, CHOP, BiP, or ERAD-related proteins SEL1L, HRD1 (aka SYVN1), DNAJC3, and DNAJB11 (**Fig 7F,G,J,K**). These findings were largely mirrored in human EndoC-βH1 cells (**Fig 7L-V**), although the immediate early response, as measured by cFOS expression, was less pronounced in EndoC-βH1 than in MIN6 β-cells. Overall, these findings support a conserved response to hypersecretion compared to ER stress in mouse and human β-cells. Not all transcriptional differences are reflected in steady-state protein abundance, or the magnitude of the alterations are not large enough to reach significance, as seen for CHOP, DNAJC3, and DNAJB11 in MIN6 (**Fig 7C,J,K**), or cFOS, PHGDH, and DNAJB11 in EndoC-βH1 (**Fig 7C,T,U**).

**Figure 7.**
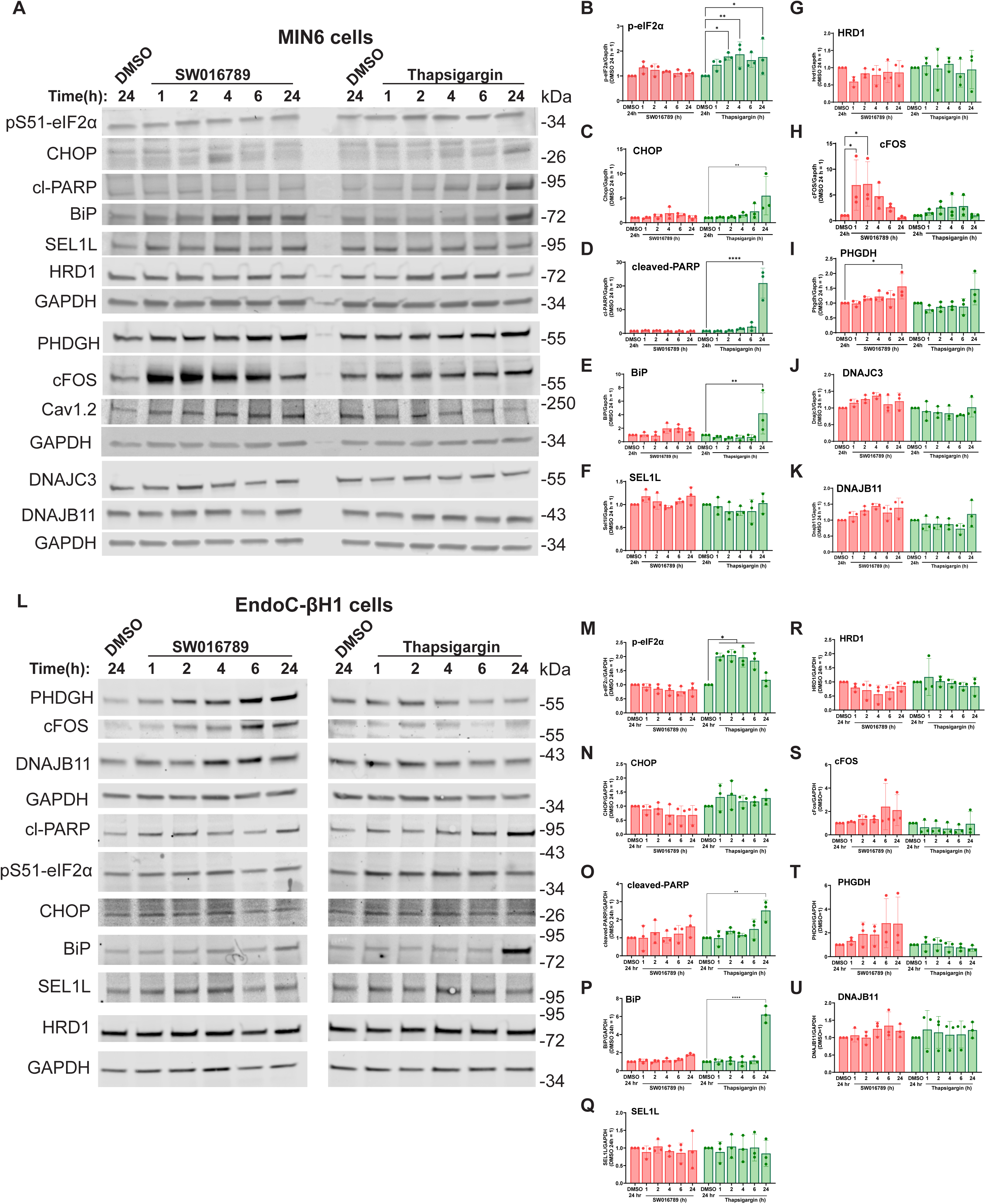
Effects of hypersecretion and ER stress on the expression of ERAD components, ER stress markers, and immediate early response genes. **A)** MIN6 β-cells were treated with SW016789 (5 μM), thapsigargin (100 nM), or DMSO (0.1%) for the indicated times and samples were analyzed by immunoblotting. **B-K)** Quantification is shown for p-eIF2α, CHOP, cleaved-PARP, BiP, SEL1L, HRD1, cFOS, PHGDH, DNAJC3, and DNAJB11, respectively. **L)** Human EndoC-βH1 cells were treated the same as in (**A**). **M-Q**) Quantification is shown for p-eIF2α, CHOP, cleaved- PARP, BiP, SEL1L, HRD1, cFOS, PHGDH, and DNAJB11, respectively. All data are the mean ± SD of N=3. *, P<0.05 vs DMSO control.

#### ERAD components HRD1, SEL1L and DERL3 are co-expressed in human islets from normal and T2D donors

In both non-diabetic and T2D pancreas tissue, HRD1 and SEL1L staining appeared across both endocrine and exocrine cells (**Fig 8A**). Within islets, HRD1 and SEL1L staining overlapped with insulin staining, indicating expression in β-cells. SEL1L and HRD1 were also present in islets from T2D donors, although islet structure appeared more disurpted in T2D compared to non-diabetic samples. HRD1 and SEL1L colocalized in islets, but the relative amount of SEL1L compared to HRD1 appeared higher in T2D islets. We reasoned that alterations in the stoichiometry of ERAD components could contribute β-cell dysfunction in T2D. Using CellProfiler, we analyzed the ratio of SEL1L to HRD1 intensity within each β-cell and plotted the distribution of the SEL1L/HRD1 ratio (**Fig 8B**). The distribution of SEL1/HRD1 ratios was significantly right-shifted in T2D (**Fig 8C**). Because the main ERAD complex contains HRD1, SEL1L, as well as DERL3, whose induction is generally more responsive to stress conditions ^39^, we also stained DERL3 in these same human samples (**Fig 8D**). DERL3 staining was enriched in islets compared to surrounding acinar tissue and in T2D, appeared to colocalize more with β-cells. DERL3 and SEL1L colocalization in islets appeared more prominent in T2D than in non-diabetic samples. Like with SEL1L/HRD1, we measured the distribution of SEL1L/DERL3 ratio in β-cells and found it was significantly left-shifted in T2D (**Fig 8E,F**). Taken together, these data suggest that at least a subset of T2D β-cells have lower HRD1 and higher DERL3 abundances relative to SEL1L. These data also confirm for the first time the co-expression HRD1, SEL1L, and DERL3 at the protein level in non-diabetic and T2D human islets. The presence of these components in T2D β-cells suggests it may be possible to target ERAD flux to ameliorate β-cell stress in diabetes.

**Figure 8.**
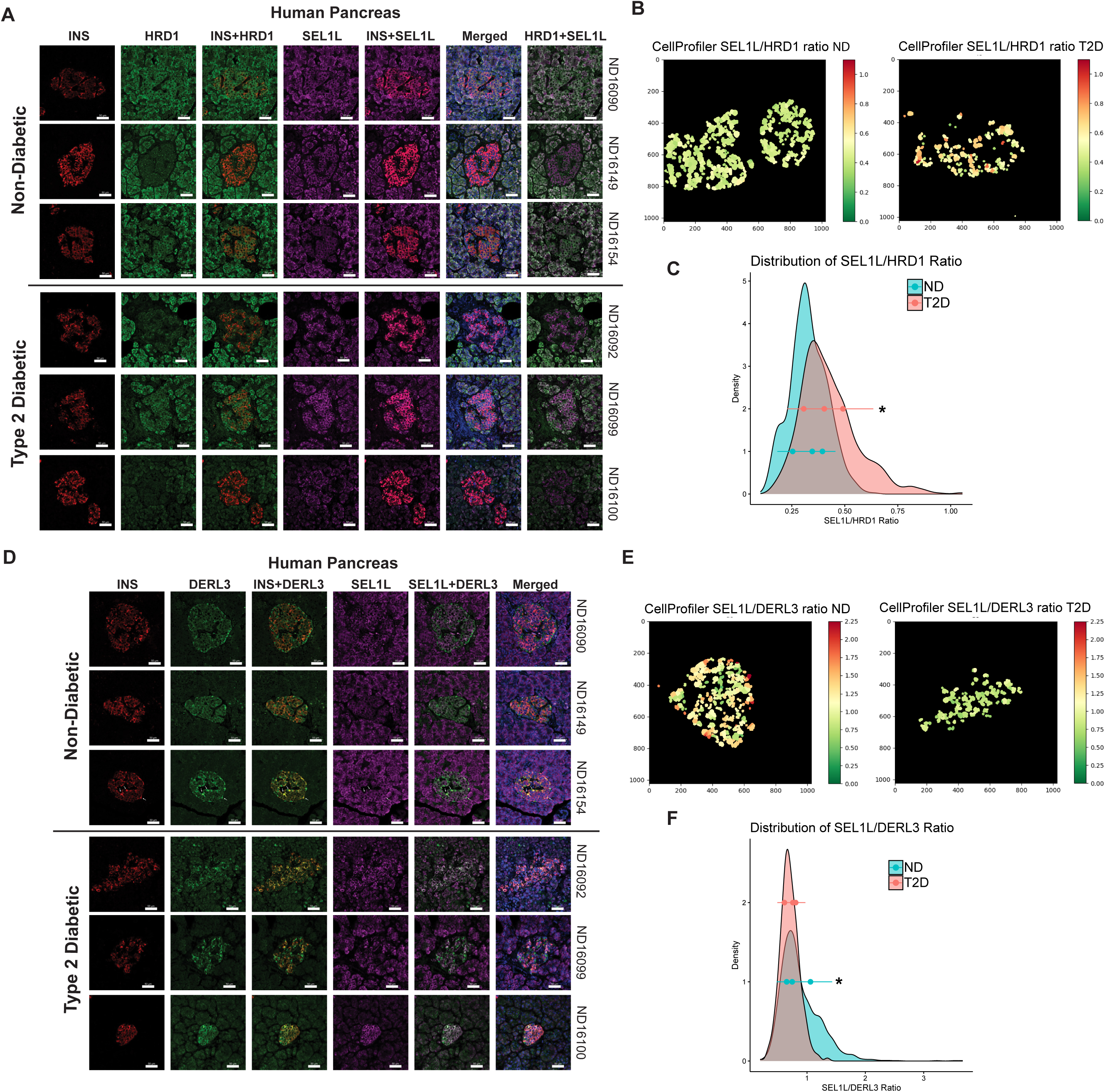
Expression of ERAD components in pancreas tissue sections from non-diabetic (ND) and type 2 diabetic (T2D) humans. **A)** Immunohistochemical staining of insulin (INS) in red, HRD1 in green, SEL1L in violet, and overlaid images showing INS+HRD1, INS+SEL1L, and HRD1+SEL1L. Donor identifiers from NDRI are shown on the right of each representative image set. **B)** CellProfiler quantification of the ratio of β-cell intensities for SEL1L and HRD1. **C)** Histogram plot of all quantified β-cell SEL1L/HRD1 ratios. *,P<0.05 by Kolmogorov-Smirnov test. **D)** Immunohistochemical staining for INS in red, DERL3 in green, SEL1L in violet, and overlaid images showing INS+DERL3 and SEL1L+DERL3. All images shown are representative of three independent ND and T2D donors. **E)** CellProfiler quantification of the ratio of β-cell intensities for SEL1L and DERL3. *,P<0.05 by Kolmogorov-Smirnov test. **F)** Histogram plot of all quantified β-cell SEL1L/DERL3 ratios. All histogram plots also show the mean ± SD of N=3 donors (3-5 imaged regions per donor).

## DISCUSSION

### The role of Vdac1 in SW016789-mediated β-cell hypersecretion

VDACs comprise three isoforms encoded by separate genes, VDAC1, VDAC2, and VDAC3. The VDACs are permeable to ions (K^+^, Na^+^, Ca^2+^) and metabolites, are expressed in β-cells, and have been pursued as possible therapeutic targets for diabetes ^40^. β-cells express VDAC1, VDAC2, and VDAC3 ^29, 41^. VDAC2 has been suggested to protect against cell death by sequestering the pro-apoptotic protein Bak ^42^.

VDAC3 may play a larger role in the testis tissue and have distinct functions from VDAC1 and VDAC2 ^43^. However, VDAC1 expression has been linked to T2D and β-cell dysfunction ^29^. VDAC1 was found to be mistargeted to the plasma membrane where it co-localized with SNARE proteins and was suggested to cause impaired GSIS in the setting of glucotoxicity and T2D ^29^. It is worth noting that mitochondrial VDAC1 protein is reduced in T2D β-cells ^44^. VDAC1 has been suggested to interact with β-cell glucokinase ^45^, which could underlie its role in dysfunctional β-cells in T2D as shown by Zhang et al. ^29^. VDAC1 could also be involved in mitochondria-ER contact sites ^46^. Indeed, VDAC1 interacts with IP3R2 and this interaction is reduced in T2D β-cells ^44^.

Our data supports that SW016789 can act through VDAC1. However, SW016789 may exhibit polypharmacology and act through additional targets. Based on our current findings and previous phenotypic characterization of SW016789 ^24^, we propose that SW016789 acts at the plasma membrane level to influence Ca^2+^ influx through VDCCs. Within this context, it is possible that some effects of SW016789 are modulated by phosphorylation of VDCCs or other ion channels that indirectly potentiate VDCC activation ^47^. While VDAC1 is purported to be present in both the mitochondrial outer membrane and the plasma membrane, our prior work with SW016789 did not uncover effects on mitochondrial function ^24^. While we have not specifically investigated plasmalemmal versus mitochondrial VDAC1, we predict that SW016789 is acting through plasma membrane-localized VDAC1. These findings are particularly relevant to the ongoing pursuit of VDAC1 as a target for inhibition in T2D and Alzheimer’s ^29, 48, 49^.

### β-cell adaptation to the spectrum of secretory stresses

Hypersecretion clearly induces a distinct response compared to pharmacological or genetic inhibition of SERCA2 (thapsigargin), a conclusion supported by our current findings and by published results of others ^50^. Hypersecretory stress is also likely different than ER stress that occurs in response to the N-linked glycosylation inhibitor tunicamycin, glucolipotoxicity, antioxidants like dithiothreitol, or induction of misfolding mutants of insulin (i.e., Ins2-C96Y; Akita mice) ^51, 52^. A major distinction is that in response to hypersecretion *in vitro*, β-cells can sacrifice their function to stave off cell death, which they cannot do in response to many other stressors. There is evidence from genetic mutations in humans (KCNQ1^53^, HNF1A^18^) and mice (SUR1^54^) that hypersecretion precedes β-cell failure and some individuals with congenital hyperinsulinism go on to develop diabetes ^55^. As noted by Nichols, York, and Remedi ^55^, pharmacological evidence suggests chronic sulfonylurea use can be detrimental to β-cell function ^21, 31, 56–59^. β-cells may deal with hypersecretion largely by downregulating insulin secretion, but over time this adaptive strategy could become maladaptive *in vivo*. Reduced insulin secretion results in hyperglycemia, and this hyperglycemia can then contribute to β-cell failure in a vicious cycle ^25^. How β-cells achieve a suppressed secretory state and the extent to which other factors may help protect β-cells from death in hypersecretion is an active area of research. ER stress responses to various insults are likely precipitated partly by the temporal dynamics of transcriptional changes and the specific genes induced. We identified multiple clusters of dynamic gene expression in our analysis that will require further investigation. For example, clusters containing immediate early genes (IEGs), or those specifically induced by thapsigargin and not hypersecretion, may contain novel cell viability regulators. For example, *Npas4* was expressed among the IEGs, and this gene has been demonstrated to have a pro-survival role in the β-cell ^60, 61^.

Other recent findings support the idea that β-cell survival under hypersecretory conditions can occur at the expense of secretory function, for example, by treating mouse islets for 48 h with the dextromethorphan metabolite dextrorphan (DXO) ^26^. This process was suggested to depend on ATF4-mediated expression of serine-linked mitochondrial one-carbon metabolism, involving the genes *Phgdh*, *Shmt2*, and *Mthfd2*. Even though we analyzed different time points, we also found genes in this pathway were induced by hypersecretion (SW_DPGP_cluster_3). With these results in mind, our findings support the possibility that the effects of SW016789-induced hypersecretion may also involve serine and one-carbon metabolism in β-cells to protect against stress-induced cell death. It is possible that hypersecretion induced by a variety of pharmacological avenues can lead to similar outcomes in the β-cell, likely via converging on enhanced Ca^2+^ influx (**Fig 9**).

**Figure 9.**
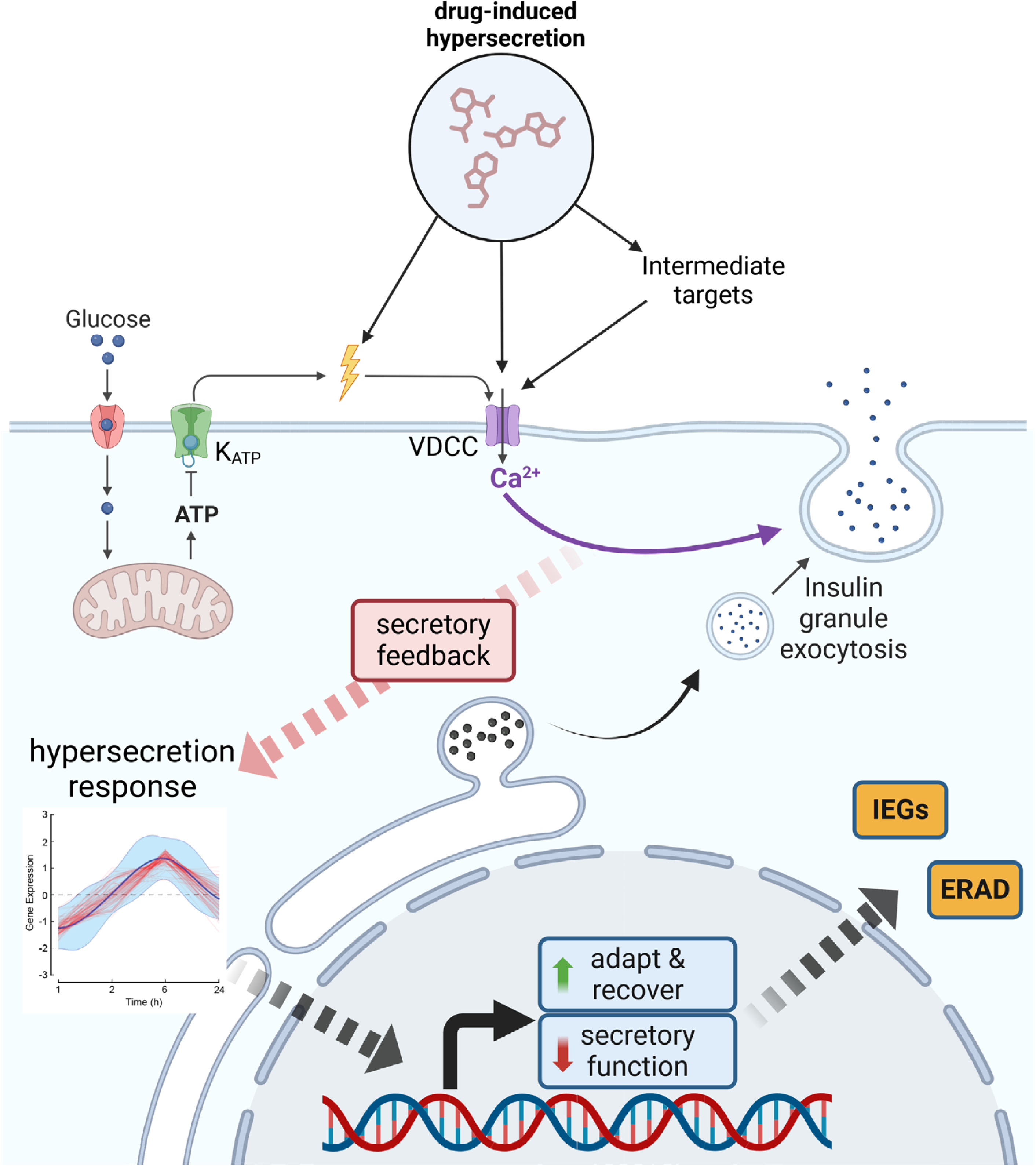
Working model of hypersecretory stress in β-cells. When β-cells are exposed to hypersecretion-inducing compounds (e.g. SW016789, sulfonylureas, DXO), regardless of their proximal protein target (e.g. VDAC1, K_ATP_, NMDAR), the key step is enhanced Ca^2+^ influx through voltage-dependent Ca^2+^ channels (VDCCs). The continuous elevation of intracellular Ca^2+^ concentration induces hypersecretion and, in parallel, activates the expression and translation of immediate early response genes within the first few hours. Subsequently, β-cells experience a negative feedback signal, which has not been elucidated, that suppresses secretion by ∼4 h. Around that same time, ER stress markers (e.g. CHOP, BiP) are transiently and mildly induced. At later time points (6-24 h), factors like PHDGH are induced, possibly because of the robust immediate early response. Critically, many gene changes detected at the mRNA level are not reflected in altered protein abundances. While the transcriptomic signature we identified is reflective of β-cells experiencing hypersecretion, the mechanisms for the disconnect between transcripts and protein abundance in these conditions are unclear.

### Blocking Ca^2+^ influx to protect against hypersecretory stress

We previously observed that nifedipine can prevent SW016789-induced hypersecretory stress and loss of function in β-cells ^24^. Furthermore, nifedipine prevents hypersecretory stress induced by all the small molecule secretion enhancers we identified in our high-throughput screen, dextrorphan (DXO), and sulfonylureas (glimepiride). This indicates that while it may be useful to understand the proximal mechanisms causing hypersecretion, Ca^2+^ influx is a key factor. Interestingly, not all inhibitors of insulin secretion confer the same protection as nifedipine. SW016789, in the presence of glucose, was able to elicit Ca^2+^ influx and insulin release even when treated with the K_ATP_ channel opener diazoxide ^24^.

Additionally, dihydropyridine-based VDCC blockers (nifedipine, nimodipine) appeared to have a superior ability to protect against SW016789 as compared to the phenylalkylamine verapamil, which required high concentrations, and diltiazem which was not protective ^24^. Data showing that VDCC activators appear to also induce hypersecretion and phenocopy SW016789 points to the importance of Ca^2+^ influx in this distinct stress response ^24, 27^. The different mechanisms of action of VDCC modulators may explain why nifedipine can protect against hypersecretory stress. Diltiazem and verapamil block ion permeation through the VDCC pore, while nifedipine interacts with a different site near the pore without blocking it ^62, 63^. Therefore, dihydropyridines can selectively inhibit VDCCs under the chronic depolarization that occurs during hypersecretion ^63, 64^.

### β-cell ERAD may be a protective mechanism upregulated during hypersecretion

Among genes induced by SW016789 in SW cluster 3, there was an enrichment of core ERAD components *Sel1l*, *Syvn1* (Hrd1), and *Derl3*. Previous work supports a requirement for well-regulated ERAD in β-cell function. HRD1 was found to be increased in T2D human islet β-cells and β-cell-specific Hrd1 overexpression in mice led to elevated fasting blood glucose levels, hyperglycemia, and impaired glucose-stimulated insulin secretion ^65^. This loss of β-cell function may be due to Hrd1-mediated degradation of MafA, a critical β-cell transcription factor. HRD1 is also upregulated in islets of Akita mice, which become diabetic because of β-cell ER stress from a misfolding insulin mutant ^66^.

However, SW016789 was previously shown not to alter MafA protein levels ^24^, nor does it significantly decrease transcript levels of β-cell transcription factors. Conversely, loss of Sel1l in mouse β-cells led to impaired function, loss of β-cell identity, and development of diabetes ^36, 37^. These findings were supported by a loss of SEL1L in β-cells from human T2D donors ^36^. For the third core ERAD component DERL1/2/3, not much is known about their regulation under normal or pathophysiological conditions in β-cells. Combined, these published findings suggest both positive and negative roles for ERAD in β-cell function and T2D pathogenesis. Our observations of HRD1 and SEL1L staining in human β-cells from normal and T2D donors suggest an imbalance in the relative abundances of HRD1 and DERL3 compared to SEL1L. Different stoichiometries between DERL3, SEL1L, and HRD1 may indicate the formation of distinct ERAD complexes that target different substrates or have different capacities. Indeed, altered levels of HRD1 can affect ERAD complex stoichiometries ^67^, and distinct ERAD complexes containing either DERL1 or DERL3 have been described ^39^. While methods to accurately monitor ERAD activity are limited, we observed decreased levels of ERAD component and substrate OS-9 after treatment with SW016789, suggesting altered ERAD activity. DERL3 may also act as a hub molecule to control the formation of different ERAD complexes. For example, DERL3 can interact with another ERAD component, HERP, and in pancreata of *Derl3* knockout mice the expression of Derl1 and Derl2 was reduced in a Herp-dependent manner ^68^. More detailed analyses and tool development are needed for ERAD flux in non-diabetic vs T2D β-cells and for understanding how hypersecretion may influence this process.

### Limitations of the study

In this work we described the identification of VDAC1 as a protein target for the small-molecule hypersecretion inducer SW016789. While this is supported by affinity purification-coupled proteomics, CETSA, RNAi, and pharmacological experiments, it is possible that SW016789 also acts through other target proteins in the β-cell. We also provided a transcriptomic resource for the time course of differential gene expression during hypersecretion induced by SW016789 or ER stress induced by thapsigargin. While these data were generated using the MIN6 mouse β-cell line, multiple validations were performed in human EndoC-βH1 cells and primary human islets. Bulk RNAseq experiments, while having the advantage of improved read depth, lack the ability to detect heterogeneity between individual β-cells during these stress responses. Single-cell transcriptomic experiments are required to better define and understand the contributions of β- cell subpopulations in hypersecretion and during diabetes pathogenesis, an issue recent studies have begun to address ^69^. Finally, the expression of ERAD components was implicated in our transcriptomic studies using SW016789. While OS-9 levels were altered, we have not exhausted all potential readouts for ERAD flux in this work. The p97/VCP inhibitor eeyarestatin I may have off-target effects ^70, 71^, and inducible β-cell knockdown/knockout of ERAD components will be ideal in the future for addressing their role(s) in adaptation to hypersecretory stress. In conclusion, further studies are needed to improve our understanding of hypersecretion downstream of enhanced Ca^2+^ influx and exocytosis and thus reveal potentially targetable components for treating diabetes or other β-cell diseases.

## METHODS

### Reagents

All chemicals and reagents are detailed in **Table S1**. The SW016789 analog, Z6292276622 (Z629), containing a fluorinated aryl azide and an alkyne-containing phenylacetylene, was synthesized by Enamine.

### Cell culture, treatments, and RNA isolation

Culture of MIN6 β-cells (<u>RRID:CVCL_0431</u>) has been described ^72^. MIN6 cells were cultured in high glucose DMEM containing 10% FBS, 50 µM beta-mercaptoethanol, 1 mM pyruvate, 100 U/ml penicillin, and 100 μg/ml streptomycin. For glucose-stimulated insulin secretion experiments, MIN6 cells were washed twice with and incubated for 2 h in freshly prepared glucose-free modified Krebs-Ringer bicarbonate (KRBH) buffer (5 mM KCl, 120 mM NaCl, 15 mM HEPES, pH 7.4, 24 mM NaHCO_3_, 1 mM MgCl_2_, 2 mM CaCl_2_, and 1 mg/ml radioimmunoassay-grade BSA). MIN6 cells were stimulated with glucose as indicated and secreted insulin and insulin content were measured using the Cisbio Insulin High Range HTRF kit (<u>AB_2890910</u>). To generate lysates for immunoblotting, cells were lysed in 1% NP40 lysis buffer (25 mM HEPES, pH 7.4, 1% Nonidet P-40, 10% glycerol, 137 mM NaCl, 1 mM EDTA, 1 mM EGTA, 50 mM sodium fluoride, 10 mM sodium pyrophosphate, 1 mM sodium orthovanadate, 1 mM phenylmethylsulfonyl fluoride, 10 μg/ml aprotinin, 1 μg/ml pepstatin, 5 μg/ml leupeptin), rotated (10 min, 4°C), and centrifuged (10,000 x g; 10 min; 4°C), or lysed in RIPA buffer (50 mM HEPES, pH 7.4, 1% Nonidet P-40, 0.5% sodium deoxycholate, 0.1% sodium dodecylsulfate, 150 mM NaCl, 1 mM EDTA, 1 mM EGTA, 50 mM sodium fluoride, 10 mM sodium pyrophosphate, 1 mM sodium orthovanadate, 1 mM phenylmethylsulfonyl fluoride, 10 μg/ml aprotinin, 1 μg/ml pepstatin, 5 μg/ml leupeptin), sonicated 5 min (30s duty cycle), and stored at -80°C. InsGLuc-MIN6 cells were generated and characterized as previously described ^72^ and were determined to be negative for mycoplasma. MIN6 β cells were treated with DMSO, SW016789 and thapsigargin for 1, 2, 6 and 24 hours and RNA was collected for each time point at the end of the time course.

### Synthesis of Z629, photoaffinity analog of SW016789

The Z629 probe was synthesized by Enamine according to the scheme shown in **Fig S1A**. <u>Step A:</u> To a solution of 2,3,4,5,6-pentafluorobenzaldehyde (13.1 g; 0.067 mol) in acetone/water (100/50 mL), sodium azide (4.57 g; 0.7 mol; 1.05 eq) was added in one portion, and the mixture was stirred at 50 °C for 16 hrs. The mixture was then cooled to room temperature (RT) and concentrated *in vacuo* below 35 °C to obtain a ∼60 mL solution. The resulting mixture was diluted with 200 mL of t-BuOMe, and the organic phase was separated and washed twice with brine, dried and concentrated to obtain crude 4-azido- 2,3,5,6-tetrafluorobenzaldehyde (13 g; 0.0593 mol, Yield = 88%, 80% purity by NMR), which was used in the next step without further purification. <u>Step B</u>: NaBH4 (2.69 g; 0.071 mol; 1.2 eq) was suspended in THF (50 mL) and the mixture cooled to –20 °C under argon atmosphere (Ar). Dimethylamine hydrochloride (5.8 g; 0.071 mol; 1.2 eq) was added portion-wise at -20 °C, and the mixture was stirred for 1 hr at this temperature and then 16 hrs at RT. The formed precipitate was filtered and the filtrate concentrated *in vacuo* to give dimethylamine borane. In a 500 mL flask 4- azido-2,3,5,6-tetrafluorobenzaldehyde (13 g; 0.059 mol; 1 eq) was dissolved in 100 mL of acetic acid, and the freshly prepared dimethylamine borane was added portion-wise at RT. The mixture was heated at 55 °C for 1hr and then concentrated *in vacuo* at 45 °C. The residue was dissolved in 200 mL of t-BuOMe, washed 4 times with 5% NaHCO_3_, dried (Na_2_SO_4_), and concentrated *in vacuo*. The residue was purified by silica gel column chromatography (eluted by Hex:EtOAc 20:1 to 1:1) to obtain (4-azido-2,3,5,6-tetrafluorophenyl)methanol (7.5 g; 0.034 mol; yield = 57%) as pale yellow solid. <u>Step C</u>: A solution of (4-azido-2,3,5,6-tetrafluorophenyl)methanol (7.3 g 0.033 mol; 1 eq) in dry dichloromethane (100 mL) was cooled in an ice bath under Ar. Pyridine (2.8 mL; 0.034 mol; 1.05 eq) and PBr_3_ (1.25 mL; 0.013 mol; 0.4 eq) were added to this stirred solution via a syringe over a period of 30 min. After 16 hrs at RT, 2-propanol (80 mL) in dichloromethane (200 mL) was added, followed after 15 min by an equal volume of 5% NaHCO_3_. Extraction with dichloromethane and evaporation yielded a light-yellow solid that was triturated with t-BuOMe. The formed precipitate was filtered and washed by t-BuOMe. The filtrate was evaporated to obtain 1-azido-4-(bromomethyl)-2,3,5,6- tetrafluorobenzene (3.4 g; 0.0119mol; Yield = 36%) as a yellow solid. <u>Step D</u>: To a solution of 1-azido-4-(bromomethyl)-2,3,5,6-tetrafluorobenzene (2.6 g; 9.15 mmol; 1 eq) in 30 mL of DMF, 2- aminobenzimidazole (1.22 g; 9.15 mmol; 1 eq) was added, followed by the addition of K_2_CO_3_ (2.02 g; 14.65 mmol; 1.6 eq). The mixture was stirred at room temperature for 16 hrs and then the mixture was poured into ice-water. The formed precipitate was filtered, washed with water, and concentrated *in vacuo* to obtain crude aminobenzimidazole **1** (2.2 g; 6.5 mmol, 60% purity by LCMS) as a light brown solid, which was used in the next step without further purification. <u>Step E</u>: The crude aminobenzimidazole **1** (2.2 g; 6.5 mmol, 60% purity by LCMS) and 2-bromo-1-(4- ethynylphenyl)ethan-1-one (1.59 g; 6.5 mmol; 1 eq) were dissolved in acetone (40 mL). Sodium iodide (1.03 g, 6.87 mmol; 1.05 eq) was added and the mixture was stirred at 50 °C for 19 hrs. The product was isolated as a solid via filtration and washed with acetone to obtain aminobenzimidazole **2** (0.8 g; 1.43 mmol; Yield = 15.6% for two steps) as an off-white solid. <u>Step F</u>: NaBH_4_ (0.024 g; 0.63 mmol; 1eq) was added portion-wise at -10 °C to a suspension of aminobenzimidazole **2** (0.35 g; 0.63 mmol; 1eq) in 10 mL of MeOH. The mixture was stirred for 30 min at room temperature and quenched with concentrated aqueous NH_4_Cl. The mixture was concentrated *in vacuo* and diluted with EtOAc and water. The organic phase was separated, dried (Na_2_SO_4_), concentrated *in vacuo* and purified by HPLC (neutral phase) to obtain Z6292276622 (Z629) (0.012 g; 0.02 mmol; Yield = 4%) as an off-white solid.

### Target deconvolution for SW016789

MIN6 β-cells and EndoC-βH1 cells are cultured in 10cm dishes in high glucose Dulbecco’s modified Eagle’s minimum essential medium (DMEM) containing 10% fetal bovine serum (FBS), 50 μM β-mercaptoethanol, 1 mM pyruvate, 100 U/mL penicillin, and 100 μg/mL streptomycin. Cells were grown to 80% confluence and then treated with 5μM Z629 and DMSO as a control for 15 minutes in the dark at room temperature. Cells are exposed to UV-B light (306 nm) for 15 minutes on ice without lid. Cells were washed with cold PBS and lysed in RIPA buffer (50 mM Hepes pH 7.4, 0.5% sodium deoxycholate, 1% NP40, 150 mM NaCl, 0.1% SDS, and Benzonase). 1000 µg lysates per reaction condition were pre-cleared by rotating with high-capacity streptavidin-agarose beads for 1 h at room temperature. Click reaction was performed with the pre- cleared lysates with 0.0125 mM biotin-azide-plus, 1.25 mM sodium ascorbate, 0.05 mM THPTA, 1.25 mM CuSO_4_ for 15 minutes in the dark at room temperature. Proteins were then precipitated in 4 volumes of acetone, and samples were centrifuged at 20000 x g for 10 minutes at 4°C to pellet insoluble proteins, then resolubilized in 4% SDS in PBS. The lysates were rotated overnight at 4°C with high-capacity streptavidin-agarose beads in affinity purification buffer (50 mM Hepes, pH 7.4; 100 mM NaCl; 1% NP-40). For SDS-PAGE beads are washed twice with both affinity purification buffer and Wash Buffer (2% SDS + 6M Urea + 150 mM NaCl) and eluted in 2% SDS, 6M Urea, 30mM Biotin, and 2M thiourea for 15 minutes at room temperature and 15 minutes at 96 degrees C. Intermediate samples were collected after pre-clearing lysates and acetone precipitation. For proteomics, beads were washed twice each with affinity purification buffer and wash buffer (2% SDS, 6M urea, 150 mM NaCl) and once with PBS to remove SDS. Beads were frozen at -80°C until processing.

### Mass spectrometry

Sample preparation, mass spectrometry analysis, bioinformatics, and data evaluation for quantitative proteomics experiments were performed in collaboration with the Indiana University Proteomics Center for Proteome Analysis at the Indiana University School of Medicine, similar to previously published protocols ^73^. *<u>Sample Preparation</u>*: On-bead samples were submitted to the IUSM Center for proteome analysis, where proteins were denatured in 8 M urea, 100 mM Tris-HCl, pH 8.5, and reduced with 5 mM tris(2-carboxyethyl)phosphine hydrochloride (TCEP, Sigma-Aldrich Cat No: C4706) for 30 minutes at room temperature. Samples were then alkylated with 10 mM chloroacetamide (CAA, Sigma Aldrich C0267) for 30 min at room temperature in the dark, prior to dilution with 50 mM Tris-HCl, pH 8.5 to a final urea concentration of 2 M for Trypsin/Lys-C based overnight protein digestion at 37 °C (0.5 µg protease, Mass Spectrometry grade, Promega V5072). *<u>Peptide Purification and Labeling</u>:* Digestions were acidified with trifluoroacetic acid (TFA, 0.5% v/v) and desalted on Pierce C18 spin columns (Thermo Fisher Cat No: 89870) with a wash of 0.5% TFA followed by elution in 70% acetonitrile 0.1% formic acid (FA). *<u>Nano-LC-MS/MS</u>:* Mass spectrometry was performed utilizing an EASY-nLC 1200 HPLC system (SCR: 014993, Thermo Fisher Scientific) coupled to Exploris 480™ mass spectrometer with FAIMSpro interface (Thermo Fisher Scientific). 1/5^th^ of each fraction was loaded onto a 25 cm EasySpray column (ES902 Thermo Fisher Scientific) at 350 nL/min. The gradient was held at 5% B for 5 minutes (Mobile phases A: 0.1% formic acid (FA), water; B: 0.1% FA, 80% Acetonitrile (Thermo Fisher Scientific Cat No: LS122500)), then increased from 4-30%B over 98 minutes; 30-80% B over 10 mins; held at 80% for 2 minutes; and dropping from 80-4% B over the final 5 min. The mass spectrometer was operated in positive ion mode, default charge state of 2, advanced peak determination on, and lock mass of 445.12003. Three FAIMS CVs were utilized (-40 CV; -55 CV; - 70CV) each with a cycle time of 1.3 s and with identical MS and MS2 parameters. Precursor scans (m/z 375-1500) were done with an orbitrap resolution of 120000, RF lens% 40, automatic maximum inject time, standard AGC target, minimum MS2 intensity threshold of 5e3, MIPS mode to peptide, including charges of 2 to 7 for fragmentation with 30 sec dynamic exclusion. MS2 scans were performed with a quadrupole isolation window of 1.6 m/z, 30% HCD CE, 15000 resolution, standard AGC target, automatic maximum IT, fixed first mass of 110 m/z. *<u>Mass spectrometry Data Analysis</u>:*

Resulting RAW files were analyzed in Proteome Discover™ 2.5 (Thermo Fisher Scientific) with either a *Mus musculus* proteome (downloaded 010917, 49922 entries) or a *Homo sapiens* reference proteome FASTA (downloaded from Uniprot 051322 with 78806 entries) plus common contaminants (73 entries) ^74^. SEQUEST HT searches were conducted with a maximum number of 3 missed cleavages; precursor mass tolerance of 10 ppm, and a fragment mass tolerance of 0.02 Da. Static modifications used for the search were carbamidomethylation on cysteine (C). Dynamic modifications included oxidation of methionine (M), deamidation of asparagine or arginine, phosphorylation on serine, threonine, or tyrosine, and acetylation, methionine loss, or methionine loss plus acetylation on protein N-termini. Percolator False Discovery Rate was set to a strict peptide spectral match FDR setting of 0.01 and a relaxed setting of 0.05. Results were loaded into Scaffold Q+S 5.2.2 (Proteome Software) for viewing. Proteomics data are provided in **Table S2**.

### Transcriptomics and data processing

Total RNA samples were first evaluated for their quantity and quality using Agilent Bioanalyzer 2100. All samples had good quality with RIN (RNA Integrity Number) greater than 9. 100 nanograms of total RNA was used for library preparation with the KAPA mRNA Hyperprep Kit (KK8581) on Biomek following the manufacturer’s protocol. Each resulting uniquely dual-indexed library was quantified and quality accessed by Qubit and Agilent TapeStation, and multiple libraries were pooled in equal molarity. The pooled libraries were sequenced with 2×100bp paired-end configuration on an Illumina NovaSeq 6000 sequencer using the v1.5 reagent kit. Samples had an average read depth of ∼52.7 million reads/sample. The sequencing reads were first quality-checked using FastQC (v.0.11.5, Babraham Bioinformatics, Cambridge, UK) for quality control. The sequence data were then mapped to either the mouse reference genome mm10, the human reference genome hg38, or the rat reference genome rn6 using the RNA-seq aligner STAR (v.2.710a) ^75^ with the following parameter: “--outSAMmapqUnique 60”. To evaluate the quality of the RNA-seq data, the number of reads that fell into different annotated regions (exonic, intronic, splicing junction, intergenic, promoter, UTR, etc.) of the reference genome was assessed using bam-stats (from NGSUtilsJ v.0.4.17) ^76^. Uniquely mapped reads were used to quantify the gene level expression employing featureCounts (subread v.2.0.3) ^77^ with the following parameters: “-s 2 -p –countReadPairs Q 10”. Transcripts per million (TPM) were calculated using length values determined by using the “makeTxDbFromGFF” and “exonsBy” functions in the “GenomicFeatures” library and the “reduce” function in the “GenomicRanges” library in R to find the length of the union of non-overlapping exons for each gene ^78^. Raw fastq, read count table, and TPMs are available on NCBI Gene Expression Omnibus (GEO) under accession number GSE194200. Read count table and TPMs are also provided as a supplemental table (**Table S3**).

### Differential expression analysis and temporal clustering by DPGP and WGCNA

Differential gene expression time course analysis was performed with edgeR. After time course analysis in edgeR, a matrix was exported containing the gene names and log_2_FC values for all genes with |log_2_FC| > 1 and adj.p.val < 0.05 for downstream analysis. *<u>DPGP:</u>* Dirichlet-Process Gaussian-Process mixture model (DPGP) was used to cluster genes according to their temporal expression patterns across the time course using an unbiased algorithm ^33^. DPGP improves upon other clustering methods like hierarchical clustering, k-means clustering, and self-organizing maps in that it does not require the user to specify the number of clusters and addresses the time series dependency issues. The number of clusters is determined by the algorithm using the input data. The resulting clusters represent different sets of response types that can be further explored to reveal novel insights into important gene regulatory pathways. DPGP runs most efficiently using a matrix of differentially expressed genes already filtered by log_2_FC and significance. To run DPGP, a Docker container was created to install DPGP and the required package versions in Python 2.7 (https://github.com/PrincetonUniversity/DP_GP_cluster). DPGP was run in this Docker environment using the following Linux commands: “DP_GP_cluster.py -i input/final_data_matrix_SW_lfc1.txt -o output/output_SW -p svg --plot –fast” and “DP_GP_cluster.py -i input/final_data_matrix_Tg_lfc1.txt - o output/output_Tg -p svg --plot –fast”. Plotted outputs from DPGP are in panels that contain transparent red lines for the expression of each individual gene within the cluster, the cluster mean, and a ribbon twice the standard deviation about the cluster means. *<u>WGCNA</u>*: Weighted-gene co-expression network analysis (WGCNA) was applied to the time course transcriptomics data ^79, 80^. A distinction between WGCNA and DPGP is that WGCNA uses the entire unprocessed read count table from the RNAseq time course. This means additional sets of genes may be discovered that did not achieve statistical significance on their own in edgeR. A set of genes all changing in the same direction, even if each individual gene is not significant, may still indicate important pathways involved in the response to SW016789 or thapsigargin. We used WGCNA parameters similar to previous applications in rodent islets ^81^. For module identification, the expression of 15,000 of the most abundant transcripts from all samples was included. A “signed” network was constructed, yielding modules where all transcripts are positively correlated, with a minimum module size of 20, and a soft thresholding power of 12. The first principal component, or module eigengene (ME) was computed for each module and used to illustrate the expression pattern across samples.

Importantly, WGCNA provides an unsupervised analysis of the correlation structure of the transcripts across all samples, with no gene annotation or sample identification information included. Gene set enrichment analysis of genes within each module was conducted using Enrichr-KG ^82^, yielding FDR-adjusted enrichment across multiple gene set libraries, including KEGG, GWAS catalog, and Gene Ontology (GO). Of the 15,000 transcripts used for module identification, ∼14,500 (∼96%) were assigned to a module, indicating a high degree of overall correlation structure among the samples. A total of 20 modules were identified, with the number of transcripts per module ranging from 3613 (turquoise) to 43 (royalblue). Among the 20 modules, 16 were significantly enriched in one or more of the Enrichr-KG libraries. WGCNA analysis results are provided in **Table S4**. To analyze the hypersecretion gene signature identified via DPGP and WGCNA, enrichment analyses, comparisons to DXO-treated mouse islet RNAseq^26^, and plot generation were completed using RStudio (R version 4.2.2) including ggplot2, ComplexHeatmap, ComplexUpset, and VennDiagram ^83^. Gene sets were analyzed with Enrichr ^84^ or Enrichr-KG ^82^ to determine enriched Biological Process GO terms. Outputs are provided in **Table S4** and **Table S5**.

#### Construction of directed graph of module relationships

To identify relationships between WGCNA co-expression modules identified from the transcriptomic sequencing of MIN6 cells, *cor.test()* in R was used to compute all pairwise correlations among the module eigengenes (ME). Only modules that enriched for one or more pathways were considered, yielding 15 modules for network construction. Network nodes represent individual modules with the top-enriched term from *Enrich-R* indicated. A full list of enriched terms for each module is provided in **Table S4**. ME-ME correlation values were used to construct a directed network in *Cytoscape*, where network edges depict correlation values that ranged from ∼|0.4| to ∼|0.9|, corresponding to P-values of ∼0.01 to 10^-15^, respectively. Edge color depicts positive *vs.* negative correlation (red, positive; blue, negative), and thickness indicates the magnitude of correlation, thicker edges showing stronger relationships.

### siRNA transfections

For siRNA experiments, MIN6 β-cells were reverse transfected. Briefly, cells were trypsinized, counted, and plated at 8e5 cells per well of a 12-well tissue culture dish.

Simultaneously, 25pmol of siRNA (ON-TARGETplus SMART pools: mouse *Vdac1* (L-047345-00- 0005), non-targeting pool (D-001810-10-20), Horizon Discovery) was complexed with Lipofectamine RNAiMax (2.5 µL) for 5 min at room temperature in serum-free DMEM in a total volume of 100 µL. Complexed siRNAs were added directly to plated cells and cultured overnight, and the medium was changed the next morning. Cells were cultured for 48 h before harvesting cell lysates for downstream analysis.

### Human pancreas tissue staining and microscopy

Formalin-fixed paraffin-embedded (FFPE) de-identified human pancreas tissue were obtained through the NDRI (**Table S6**). Tissue was processed into 5 µm sections and mounted on glass slides at the Indiana University School of Medicine Histology Lab Service Core. Slides were deparaffinized by xylene and ethanol washes. Antigen retrieval was performed by heating for 40 min in an Epitope Retrival Steamer with slides submerged in Epitope Retrieval Solution (IHC-Tek). Subsequently, slides were placed onto disposable immunostaining coverplates and inserted into the Sequenza slide rack (Ted Pella / EMS) for washing, blocking, and antibody incubations. After three 10 min washes in IHC wash buffer (0.1% Triton X-100 and 0.01% sodium azide in PBS pH 7.4), slides were blocked for 1 h at room temperature in normal donkey serum (NDS) block solution (5% donkey serum, 1% bovine serum albumin, 0.3% Triton X-100, 0.05% Tween-20, and 0.05% sodium azide in PBS pH 7.4). Slides were then incubated overnight at 4°C with primary antibodies (**Table S1**) diluted in NDS block solution.

After three washes in IHC wash buffer 200 µL each), slides were incubated in secondary antibodies in NDS block solution for 1 h at room temperature. The washed slides were mounted in polyvinyl alcohol (PVA) mounting medium (5% PVA, 10% glycerol, 50mM Tris pH 9.0, 0.01% sodium azide) with DAPI (300 nM) added and imaged on Zeiss LSM710 confocal microscope equipped with a Plan-Apochromat 20x/0.8 objective (#420650-9901). Images were processed in the Zeiss Zen software to add scale bars, set coloration for channels, and generate merged images. Scale bars indicate 50 µm. Intensities of SEL1L, HRD1, and DERL3 staining within individual β-cells, and the SEL1L/HRD1 and SEL1L/DERL3 ratio calculations were performed using CellProfiler ^85^. Data were processed further in R to generate distribution plots.

### Human islet culture and treatments

Cadaveric human islets were obtained through the Integrated Islet Distribution Program (IIDP) and Prodo Labs. Islets were isolated by the affiliated islet isolation center and cultured in PIM medium (PIM-R001GMP, Prodo Labs) supplemented with glutamine/glutathione (PIM-G001GMP, Prodo Labs), 5% Human AB serum (100512, Gemini Bio Products), and ciprofloxacin (61-277RG, Cellgro, Inc) at 37°C and 5% CO2 until shipping at 4°C overnight. Human islets were cultured upon receipt in complete CMRL-1066 (containing 1 g/L (5.5 mM) glucose, 10% FBS, 100 U/ml penicillin, 100 μg/ml streptomycin, 292 µg/mL L-glutamine). For drug treatments, human islets (50-75) were hand-picked under a dissection microscope and transferred to low-binding 1.5 ml tubes (50 islets/tube). Islets were cultured 24 h in 500 µL of complete CMRL-1066 medium containing indicated drug treatments. Islets were harvested for RNA isolation using the Quick-RNA Microprep (Zymo). Human islet characteristics and donor information are listed in **Table S7**.

### Relative gene expression measurements by qPCR

RNA was isolated using the Aurum Total RNA Mini kit (Bio-Rad). Briefly, after indicated treatments, medium was removed from cells and lysis buffer (with β-mercaptoethanol) was added to cells. Cells were scraped, lysates transferred to 1.5 mL tubes on ice and then transferred to -80°C for storage. Samples were processed according to the kit manufacturer’s instructions, including on-column digestion with RNase-free DNase. RNA concentration was measured using an Implen Nanophotometer or Nanodrop spectrophotometer and verified to have A_260/280_ ratios >2.0. 1000 ng of RNA was converted into cDNA using the iScript cDNA synthesis kit (Bio-Rad) following manufacturer instructions and the resulting cDNA was diluted 20-fold with water. One μl of diluted cDNA was used in 10 μl qPCR reactions using 2X SYBR Bio-Rad master mix and 250 nM of each primer. Reactions were run in 384-well format on a CFX384 (Bio-Rad) or QuantStudio 5 (Thermo). qPCR data was analyzed using CFX Manager (Bio-Rad) with 18S RNA as the reference gene. Relative expression was calculated by the 2^-ΔΔCt^ method. All primer details are listed in **Table S1**.

### Cellular thermal shift assays (CETSA)

MIN6 cells were cultured in 10cm dishes and pre-treated with either media containing DMSO or 10μM SW016789 for 30 minutes. The cells are trypsinized, counted, spun down and resuspended in 500μl PBS containing either DMSO or 10μM SW016789. 50μl cell suspension is aliquoted into a PCR strip tube, incubated for 5 min at RT, and heated in a thermocycler for 3 minutes. After cooling down, 50μl 2x lysis buffer (0.5% Triton-X100, 20% Glycerol, 274 mM NaCl, 50 mM Hepes pH 7.4, 2mM NaVO4, 2mM EGTA, 2mM EDTA, 20mM NAPP, 100mM NaF) is added to each tube of the PCR strip and transferred to a 1.5ml microcentrifuge tube and placed on ice for 20 minutes for cell lysis to occur. The tubes are centrifuged at 20,000*g* at 4ᵒC for 20 minutes to pellet out the aggregated insoluble material.

### Cell stress assays

To measure cells going through apoptosis we measured caspase activity with the Caspase3/7-Glo assay (Promega). The Sartorius Incucyte S3 was used to measure the induction of ER stress over time in EndoC-βH1. For this, we transduced cells with a BacMam system expressing SantakaRed constitutively and mNeonGreen under the control of IRE1-mediated intron splicing according to the manufacturer instructions (Montana Molecular).

### SDS-PAGE and immunoblotting

40-50 µg of cleared cell lysates were separated on 4-20% gradient gels (Mini-Protean or Criterion TGX, Bio-Rad) by SDS-PAGE and transferred to nitrocellulose for immunoblotting. All membranes were blocked in Odyssey blocking buffer (Licor) diluted in TBS-T (20 mM Tris-HCl pH 7.6, 150 mM NaCl, 0.1% Tween-20) for 1 h before overnight incubation with primary antibodies diluted in blocking buffer. Complete antibody information is available in **Table S1**. After three 10 min washes in TBS-T, membranes were incubated with fluorescent secondary antibodies 1 h at room temperature. After three 10 min washes in TBS-T, membranes were imaged on a Licor Odyssey scanner.

### Secreted Insulin-Gaussia luciferase assays

InsGLuc secretion assays were performed as previously described ^72, 86^. Briefly, InsGLuc-MIN6 cells were plated in 96-well dishes at 1e5 cells/well and cultured for 3-4 days before compound treatment either overnight (24 h) in medium or acutely (1 h) in KRBH. Treatments in 96-well assays were performed with at least three replicate wells and in at least three independent passages of cells. For assays, cells were washed twice with KRBH and preincubated in 100 µl of KRBH (250 µM diazoxide where indicated) for 1 h. The solution was then removed, and cells were washed once with 100 µl of KRBH and then incubated in KRBH with or without the indicated stimulation and compound treatments for 1 h. 50 µl of KRBH was collected from each well and pipetted into a white opaque 96-well assay plates. Fresh GLuc assay working solution was then prepared by adding coelenterazine stock solution into assay buffer to a final concentration of 10 µM. 50 µl of working solution was then rapidly added to the wells using an Integra Voyager 1250 µl electric multi-channel pipette for a final concentration of 5 µM coelenterazine. Plates were spun briefly, and luminescence was measured on a Synergy H1M2 plate reader (BioTek). The Gen5 software protocol was set to shake the plate orbitally for 3 seconds and then read the luminescence of each well (integration, 100 ms; gain,150). High-throughput screening of compounds in the InsGLuc-MIN6 β-cell line was previously described ^24, 32^.

### Statistical Analysis

Quantitated data are expressed as mean ± SD. Data were evaluated using one-way or two-way ANOVA as indicated, or Kolmogorov–Smirnov tests for distribution plots, and considered significant if P < 0.05. Graphs were made in GraphPad Prism 9, and figures were assembled in Adobe Illustrator. Model figures were created with BioRender.

## Author Contributions

GR – performed experiments, analyzed data, investigation, software, visualization, and writing – review & editing; AO – performed experiments, analyzed data, and visualization; KRS – investigation, formal analysis; DDB – investigation; GK – software, investigation; MBK, NM, ATT, and TR – methodology; ZT – investigation, formal analysis; EHD – investigation, formal analysis, writing – review & editing, data curation; ALM – supervision; CHM – software; MPK – software, methodology, data analysis, visualization; ADA – supervision, funding acquisition; TSJ – software, methodology, supervision, validation; MAK – conceptualization, funding acquisition, project administration, supervision, writing – review & editing, data curation, investigation, software, methodology, formal analysis.

## Acknowledgments & Funding

We would like to thank colleagues at the Indiana Biosciences Research Institute for helpful discussions, including Jonathan Flak and Erica Cai. Thanks to Jessica Maiers at IUSM for sharing the OS-9 antibody. Thanks to Pathum Weerawarna at IUSM for UV crosslinking and click chemistry advice. This work was supported by a Pilot and Feasibility Award (to M.A.K.) within the CDMD NIH/NIDDK Grant Number P30 DK097512. The proteomics work was supported, in part, by the Indiana Clinical and Translational Sciences Institute (funded in part by Award Number UL1TR002529 from the National Institutes of Health, National Center for Advancing Translational Sciences, Clinical and Translational Sciences Award) and, in part, by the IU Simon Comprehensive Cancer Center Support Grant (Award Number P30CA082709 from the National Cancer Institute).

Acquisition of the IUSM Proteomics core instrumentation used for this project was provided by the Indiana University Precision Health Initiative. Sequencing analysis was carried out in the Center for Medical Genomics at Indiana University School of Medicine, which is partially supported by the Indiana University Grand Challenges Precision Health Initiative. We thank personnel Ed Simpson, Hongyu Gao, Fang Fang, and Maks Luthra in both the Center for Computational Biology and Bioinformatics and the Center for Medical Genomics, directed by Dr. Yunlong Liu.

Immunohistochemistry slide preparation was supported by the Histology Core of the Indiana Center for Musculoskeletal Health at IU School of Medicine and the Indiana Clinical Translational Sciences Institute (CTSI). Human pancreatic islets and/or other resources were provided by the NIDDK-funded Integrated Islet Distribution Program (IIDP) (RRID:SCR_014387) at City of Hope, NIH Grant # 2UC4DK098085. This work was also supported in part by funding from the U.S. Department of Veterans Affairs (IK2 BX004659 to ATT), T32 DK064466 to Indiana University Diabetes and Obesity Research Training Program (NM), AnalytixIN (TSJ), Indiana University Precision Health Initiative (TSJ), 1R01GM148970 (TSJ), 1R21CA264339 (TSJ), the National Institute on Aging of the National Institutes of Health (NIH) under award U54AG065181 (TR), and R01DK101573, R01DK102948, and RC2DK125961) (to ADA), the University of Wisconsin-Madison, Department of Biochemistry and Office of the Vice Chancellor for Research and Graduate Education with funding from the Wisconsin Alumni Research Foundation (to MPK), and by the American Diabetes Association (grant no.: 7-21- PDF-157) (to CHE).

## Supplementary Material

Supplementary figures and tables are included as part of this manuscript.

## Data Availability Statement

The datasets generated for this study can be found in the NCBI Gene Expression Omnibus GSE194200 and ProteomeXchange Partner MassIVE under repository ID MSV000094310.

---

**Figure S1.**
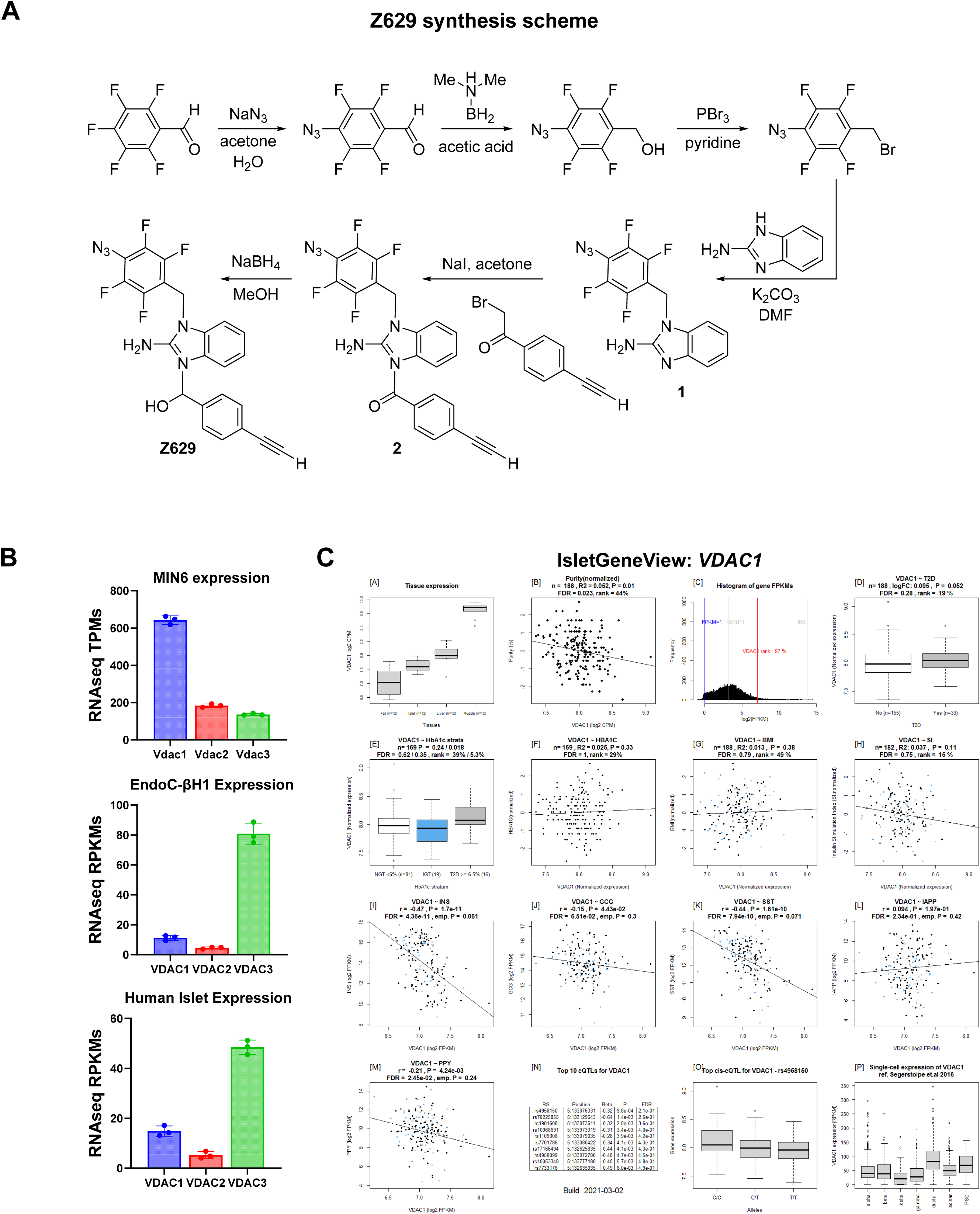
SW016789 photoaffinity probe synthesis and β-cell VDAC expression. **A)** Synthetic scheme for Z6292276622 (Z629) which incorporates an aryl azide for UV crosslinking and an alkyne for click chemistry. **B)** Expression of *VDAC1*, *VDAC2*, and *VDAC3* in MIN6, EndoC-βH1, and human islets. MIN6 TPMs are from our own RNAseq data (GSE194200). EndoC-βH1 and human islet RPKM data is from Fred RG, et al.^87^. **C)** IsletGeneView report on VDAC1 expression in a large human islet RNAseq dataset containing non-diabetic and type 2 diabetic donor islets. VDAC1 is well-expressed in islets and is negatively correlated with *INS* expression.

**Figure S2.**
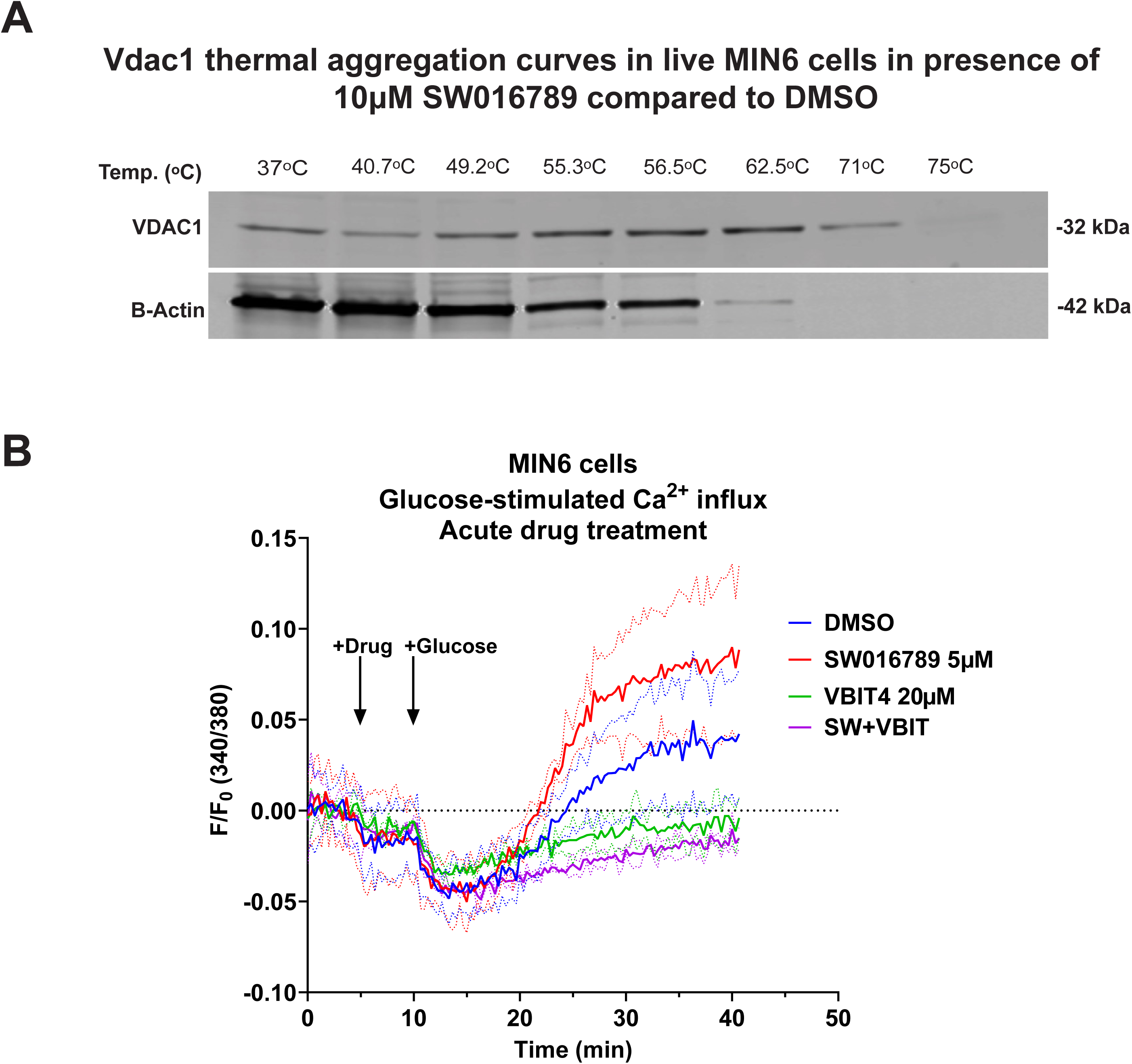
VDAC1 thermal stability and impact of VBIT4 on Ca^2+^ influx. **A)** Thermal stability of endogenous VDAC1 (37°C - 75°C) in the absence of ligand shows stability up to 71°C compared to β-actin in MIN6 β-cells. **B)** Glucose-stimulated Ca^2+^ influx in MIN6 β-cells treated acutely with SW016789 (SW, 5 μM), VBIT4 (20 μM), or both.

**Figure S3.**
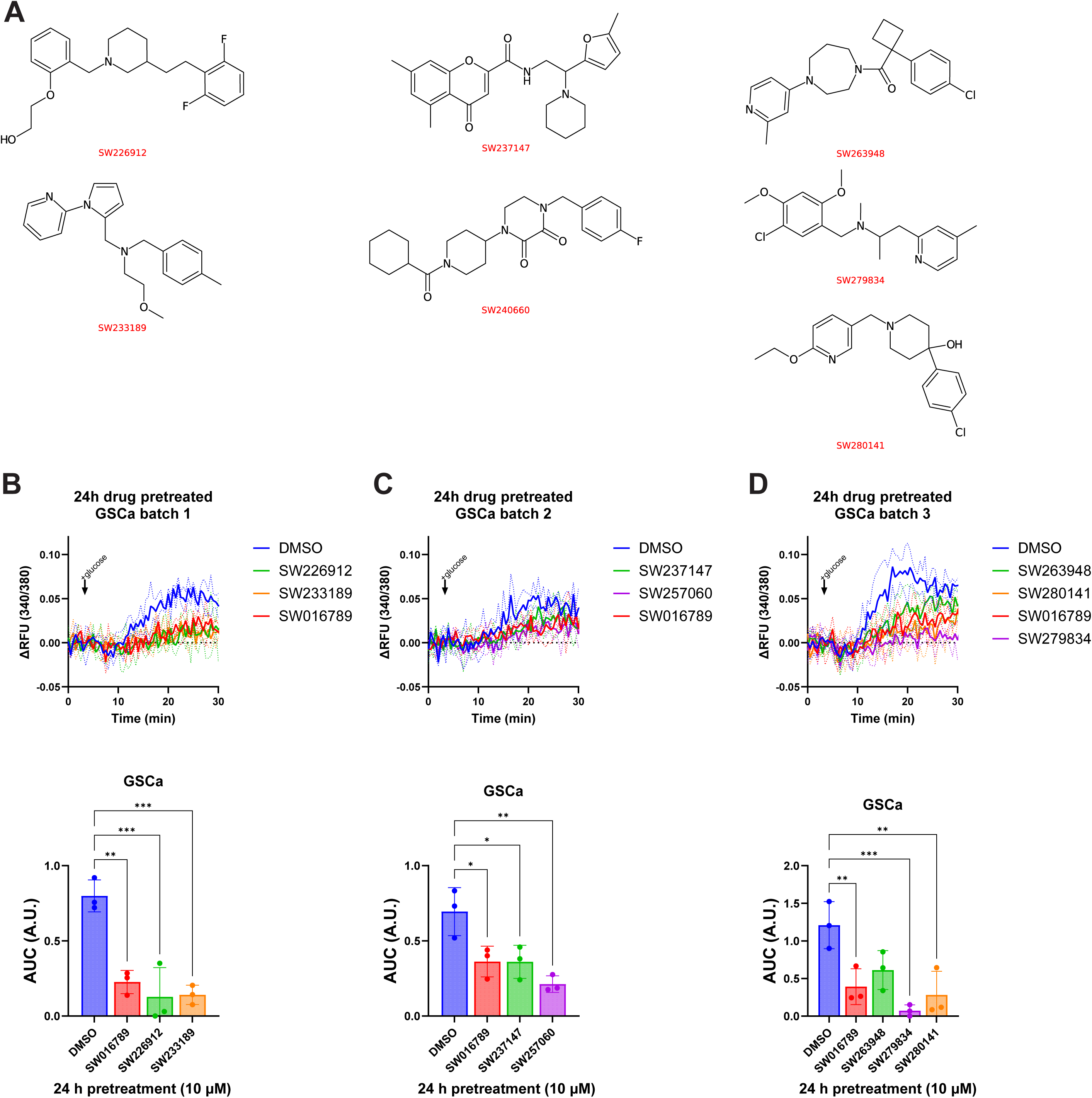
Diverse small molecule hypersecretion inducers phenocopy SW016789. **A)** Structures are shown for high-throughput screening hits previously identified ^32^. **B)** Glucose-stimulated Ca^2+^ influx after 24 h pretreatment with compounds (10 μM) from (A) or SW016789 (5 μM) as a positive control. Data are the mean ± SD of N=3 experiments. *, P<0.05.

**Figure S4.**
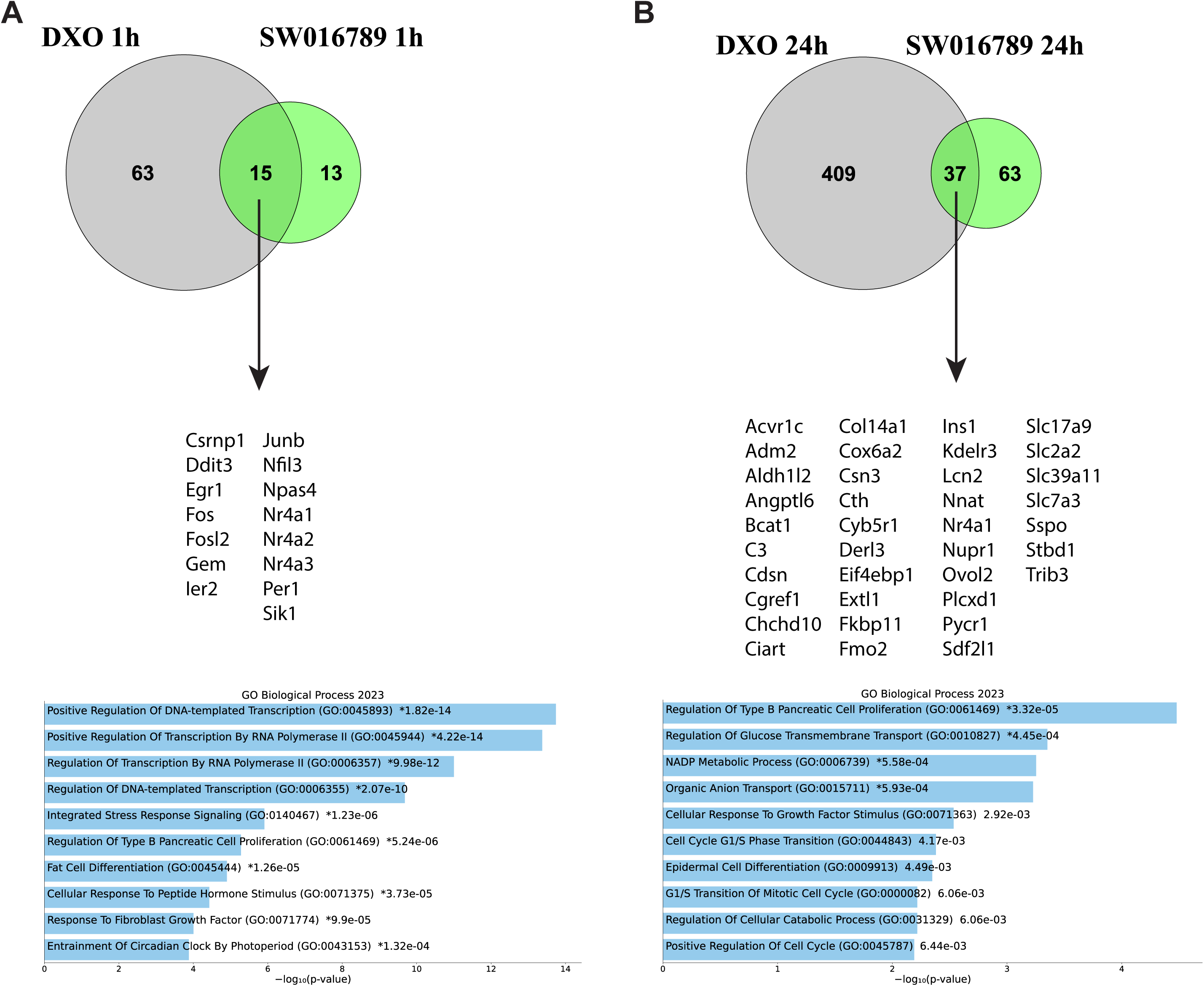
Differential gene expression comparison of SW016789 and DXO. Published transcriptomics data of DXO-treated mouse islets ^26^ was compared to our SW016789-treated mouse MIN6 β-cell transcriptomics data at the corresponding 1 h (**A**) and 24 h (**B**) time points. Below the venn diagrams the overlapping genes are shown (also in Table S5) as well as the GO Biological Process enrichment.

**Figure S5.**
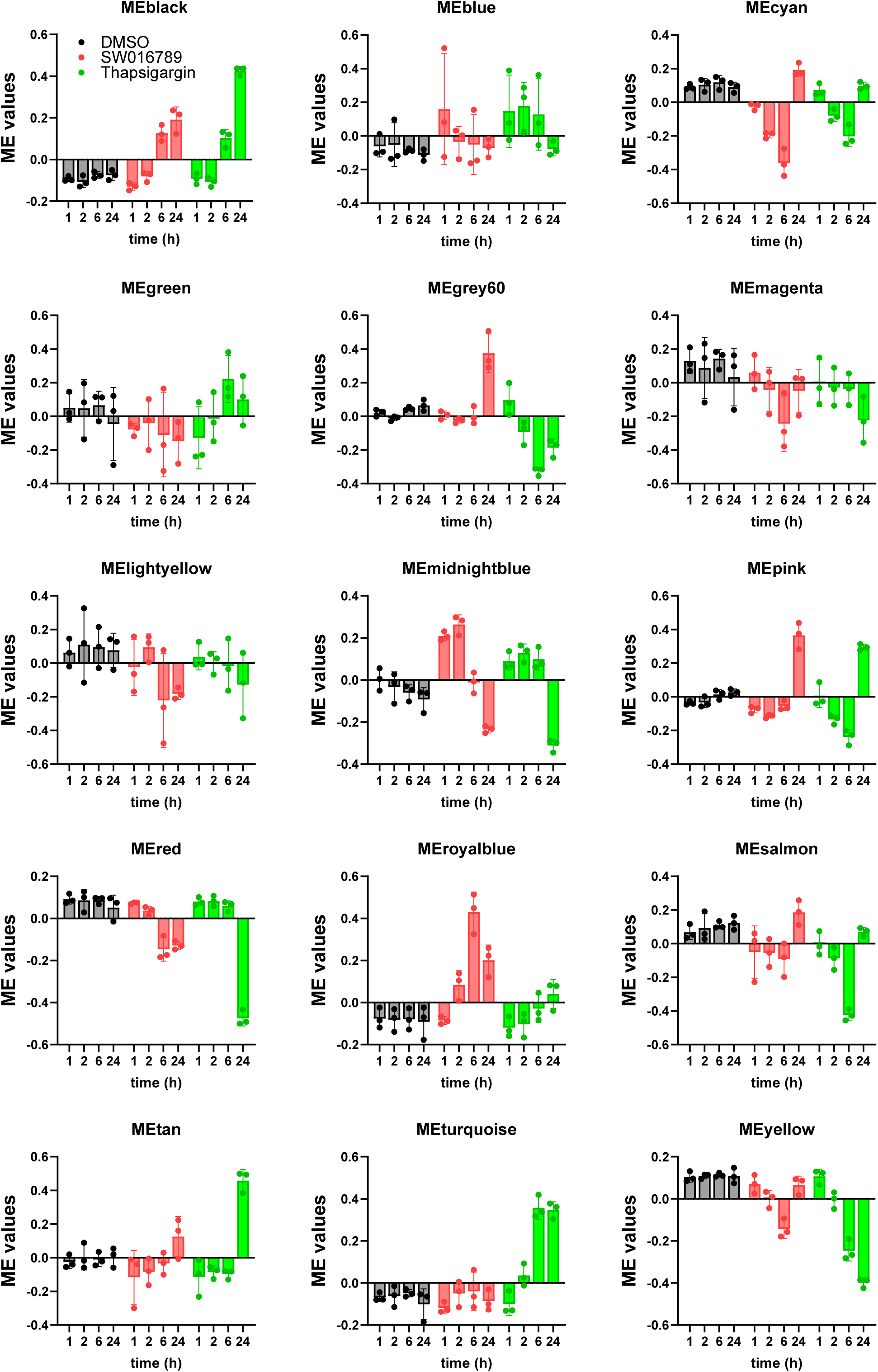
WGCNA modules identified from hypersecretion and ER stress time course transcriptomics. Bar graphs show the mean ± SD of module eigengene (ME) values for each treatment within each module across the time course.

