## Supplementary material for "Target deconvolution of an insulin hypersecretion-inducer acting through VDAC1 with a distinct transcriptomic signature in beta-cells": Table S7

### Checklist for reporting human islet preparations used in research

Adapted from Hart NJ, Powers AC (2018) Progress, challenges, and suggestions for using human islets to understand islet biology and human diabetes. Diabetologia https://doi.org/10.1007/s00125-018-4772-2

| Islet preparation | 1 | 2 | 3 | 4 | 5 | 6 | 7 | 8[a] |
|---|---|---|---|---|---|---|---|---|
| **MANDATORY INFORMATION** | | | | | | | | |
| Unique identifier | SAMN30986138 | HP-22278-01 | HP-23153-01 | HP-23166-01 | SAMN34033792 | | | |
| Donor age (years) | 67 | 69 | 51 | 58 | 39 | | | |
| Donor sex (M/F) | male | male | Female | Female | Male | | | |
| Donor BMI (kg/m$^2$) | 34.4 | 29.1 | 26.4 | 28.8 | 33.4 | | | |
| Donor $HbA_{1c}$ or other measure of blood glucose control | 5.3% | 5.4% | 5.3% | 5.6% | 5.0% | | | |
| Origin/source of islets[b] | | | | | | | | |
| Islet isolation centre | Scharp-Lacy (Prodo) | Prodo Labs | Prodo Labs | Prodo Labs | Southern California Islet Cell Resource Center | | | |
| Donor history of diabetes? Please select yes/no from drop down list | No | No | No | No | No | | | |
| **If Yes, complete the next two lines if this information is available** | | | | | | | | |
| Diabetes duration (years) | | | | | | | | |
| Glucose-lowering therapy at time of death[c] | | | | | | | | |
| **RECOMMENDED INFORMATION** | | | | | | | | |
| Donor cause of death | head trauma | anoxia | stroke | stroke | head trauma | | | |

| | | | | | |
|---|---|---|---|---|---|
| Warm ischaemia time (h) | | | | | |
| Cold ischaemia time (h) | 10h 5m | | | | 12h 41m |
| Estimated purity (%) | 90 | 95 | 85 | 80-85 | 80 |
| Estimated viability (%) | 95 | 95 | 95 | 95 | 97 |
| Total culture time (h)[d] | | | | | |
| Glucose-stimulated insulin secretion (static culture by IIDP)[e] | 4.6 | | | | 2.2 |
| Handpicked to purity? Please select yes/no from drop down list | Yes | Yes | Yes | Yes | Yes |
| Additional notes: Related to Figure: | Fig 1B | Fig 1B | Fig 1B | Fig 1B | Fig 1B |
| Additional notes -Culture time prior to shipment | 3d 20h | | | | 2d 7h |

[a]If you have used more than eight islet preparations, please complete additional forms as necessary
[b]For example, IIDP, ECIT, Alberta IsletCore
[c]Please specify the therapy/therapies
[d]Time of islet culture at the isolation centre, during shipment and at the receiving laboratory
[e]Please specify the test and the results
